# From messy chemistry to ecology: autocatalysis and heritability in prebiotically plausible chemical systems

**DOI:** 10.1101/2024.08.03.606486

**Authors:** Tymofii Sokolskyi, Bruno Cuevas Zuviria, Sydney Gargulak, Esau Allen, David Baum

**Affiliations:** Wisconsin Institute for Discovery, University of Wisconsin-Madison, Madison, WI 53715, USA; Department of Botany, University of Wisconsin-Madison, Madison, WI 53706, USA; Centro de Biotecnología y Genómica de Plantas, Universidad Politécnica de Madrid (UPM) - Consejo Superior de Investigaciones Científicas (CSIC-INIA), Pozuelo de Alarcón, Madrid, Spain 28223

**Keywords:** origin of life, autocatalysis, pyrite, heritability, computational chemistry, LCMS

## Abstract

A key question in origins-of-life research, is whether heritability, and thus evolution, could have preceded genes. Out-of-equilibrium chemical reaction networks with multiple autocatalytic motifs may provide chemical “memory” and serve as units of heritability, but experimental validation is lacking. We established conditions that may be conducive to the emergence of heritable variation and developed methods to search for heritability and autocatalysis. We prepared a food set (FS) of three organic species, three inorganic salts and pyrite. We conducted a serial dilution experiment where FS was incubated for 24 hours, after which a 20% fraction was transferred into freshly prepared FS that went through the same procedure, repeated for 10 generations. To serve as controls, we also incubated the fresh solutions in each generation. We compared the chemical composition of transfer vials and no-transfer controls using liquid chromatography-mass spectrometry (LCMS), with metrics adapted from ecology and evolutionary biology. While variability was high, focusing on a subset of chemicals with more consistent patterns revealed evidence of heritable variation among vials. Using rule-based chemical reaction network inference, constrained by the LCMS data, we identified a plausible FS-driven chemical reaction network that was found to contain numerous autocatalytic cycles.

## Background

The capacity for evolution is one of the key attributes of life (Joyce, 1994). As a result, finding chemical systems that are capable of adaptive evolution, yet simple enough to emerge spontaneously under prebiotic conditions is a central goal of origins-of-life research (Paul & Joyce, 2004). A necessary prerequisite for adaptive evolution is a mechanism to generate heritable variation. In modern evolutionary biology, heritability is due to the presence of variation in self-replicating genetic molecules, which tends to result in a correlation between the traits of parents and their progeny. However, theoretical work suggests that heritable variation need not depend upon the existence of genetic molecules like RNA or DNA but can also arise in chemical reaction networks (CRNs) that contain self-amplifying, which is to say autocatalytic, motifs (Wachtershauser 1988; Segre et al., 2001a; Vasas et al. 2012; Baum 2018). However, to date, experimental data validating these claims are lacking.

It has been proposed that life emerged when CRNs driven out of equilibrium by a continuing influx of “food” established some form of spatial structure, for example in micelles (Segre et al., 2001a) or mineral surfaces (Wachtershauser et al. 1988; Baum, 2018). Given occasional seeding of new autocatalytic motifs by rare seeding events, different spatial locations could establish different dynamical states that would tend to be passed on to other locations via dispersal (Peng et al., 2022; Baum et al. 2023). However, such autocatalytically “encoded” heritability has yet to be shown in the lab and, indeed, evidence of autocatalysis in small-molecule organic chemistry remains scant (Orgel, 2008).

Since heritability depends on their being multiple autocatalytic motifs (Vasas et al., 2015), a minimal requirement for a CRN to show evolvability is that it generates a diverse set of chemical compounds. Such compositional diversity is readily achieved in prebiotic chemistry via combinatorial explosions (Warr, 1997; Colon-Santos et al., 2019; Matange et al., 2023), especially in the presence of heat and catalytic surfaces (de Graaf et al., 2023). It seems theoretically possible, therefore, to initiate combinatorial explosions in the lab in such a way as to foster sufficient autocatalytic complexity to result in the emergence of heritable variation.

Despite the desirability of creating laboratory conditions suitable for the emergence of heritability and evolution, there remain many challenges. The simplest to solve is the establishment of systems that can be interpreted as “parents” and “offspring” (Baum & Vetsigian, 2017). This could be achieved using a set of continuously stirred tank reactors (Happel & Stadler, 1999) or by establishing multiple parallel lineages in a recursive transfer paradigm, also known as chemical ecosystem selection (Vincent et al., 2019).

Multiple parent-offspring lineages can be used to calculate an analog of narrow-sense heritability, by looking at the slope of a linear regression of parent-offspring chemical characteristics (Sokolskyi et al., 2024). Other options include an approach developed by Guttenberg et al. (2015), which uses principal component analysis to estimate the number of heritable states in a complex system, or a method proposed by Segrè et al. (2001b) based on the threshold cosine similarity among vectors of chemical composition.

In this paper we used a chemical ecosystem selection paradigm in conditions where a simple, small-molecule food solution yields a combinatorial explosion of chemical complexity. Then, using the detailed chemical fingerprints generated by liquid-chromatography/mass-spectrometry (LCMS), we evaluated evidence of heritability. We find that the vials receiving transfers from prior generations tend to have more chemical diversity than no-transfer controls and that, in many generations, the parent and offspring vials tend to be more chemically similar to each other than to other vials. Finally, *in silico* chemistry analyses, guided by the LCMS data, revealed evidence of autocatalytic processes that could plausibly explain heritability. Our analysis not only provides the first direct (if weak) evidence of heritability but also suggests promising avenues for further research into the de novo emergence of heritability and evolutionary change.

## Materials & Methods

Solution preparation. We prepared a food set (FS) solution consisting of 6 compounds and a mineral, pyrite (**Table 1**), which is known for its catalytic activity (de Graaf et al., 2023). All components except pyrite were mixed in water and stored at 4°C until use. An aliquot of FS was added to pyrite every generation of the experiment. The organic components, formic acid, acetic acid and methanol, are prebiotically plausible, and were likely abundant on early Earth (McColom & Seewald, 2007; Mohammadi et al., 2020). Bicarbonate may support pyrite-catalyzed carbon fixation, especially under high temperature and pressure (Cody et al., 2004). Ammonium hydroxide and trimetaphosphate serve as sources of nitrogen and phosphorus respectively.

**Table 1.**
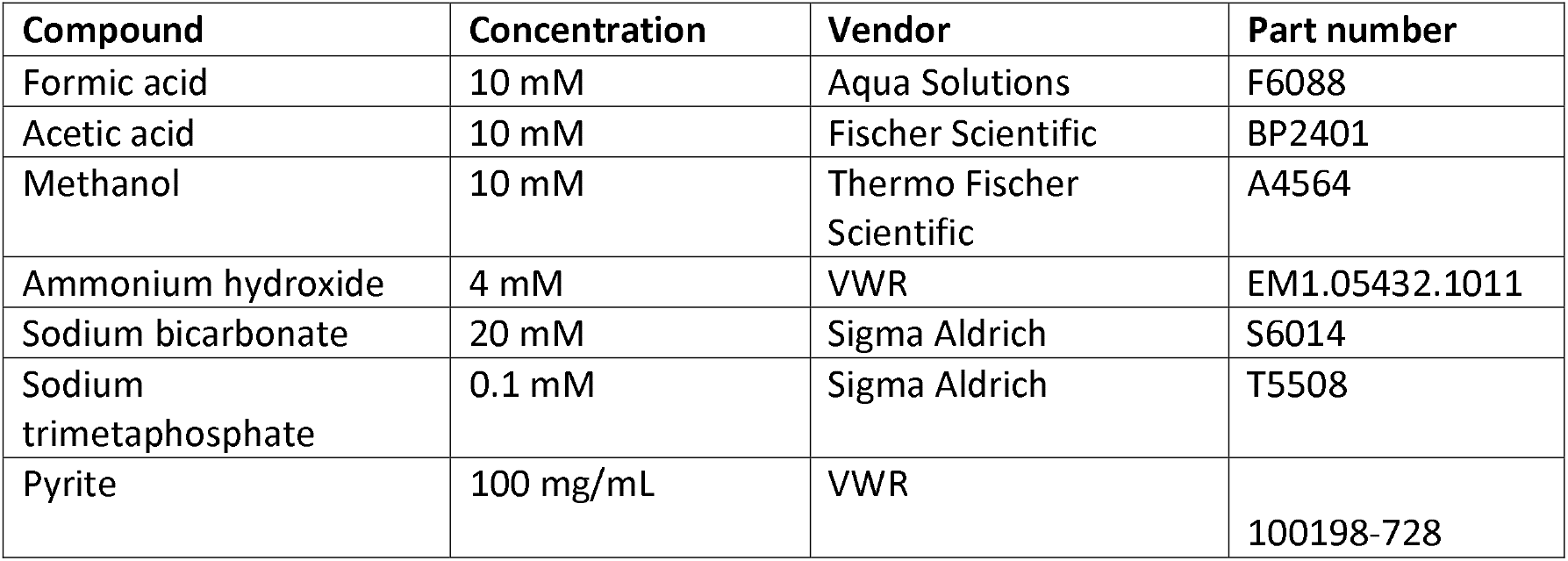
Composition of the food set (FS).

Experimental design. We conducted chemical ecosystem selection involving a transfer-with-dilution protocol (**Fig. 1**). First, we aliquoted 2 mL of FS with pyrite into glass vials (Neta Scientific, RES-24658), sealed with rubber stoppers (Neta Scientific, W224100-400) and aluminum seals (Neta Scientific, 224177-05). This procedure was conducted in an anaerobic chamber (95% N2, 5% H2). The starting vials, designated generation 0, were autoclaved using a liquid cycle (30 mins, 121°C, 30 psi) and incubated for 24 hours in a 40°C orbital shaking incubator. Generation 1 vials received 1.6 mL FS and ∼200 mg pyrite and either 0.4 mL (20%) of the solution from a “parental” generation 0 vial or an additional 0.4 mL of fresh FS. Vials receiving transfers (TRs) and no transfer controls (NTCs) were again autoclaved and incubated for 24 hours. This procedure was repeated for 10 generations, with vials from each generation stored at -80°C until analysis. In this work, our main analytical focus was to compare TRs to NTCs every generation, as it allows us to control for experimental and analytical artifacts. We conducted two identical replicate experiments, each with 10 TR lineages and 10 NTC pseudo-lineages per generation. The results shown in this study are based on a combined analysis of both experiments.

**Figure 1.**
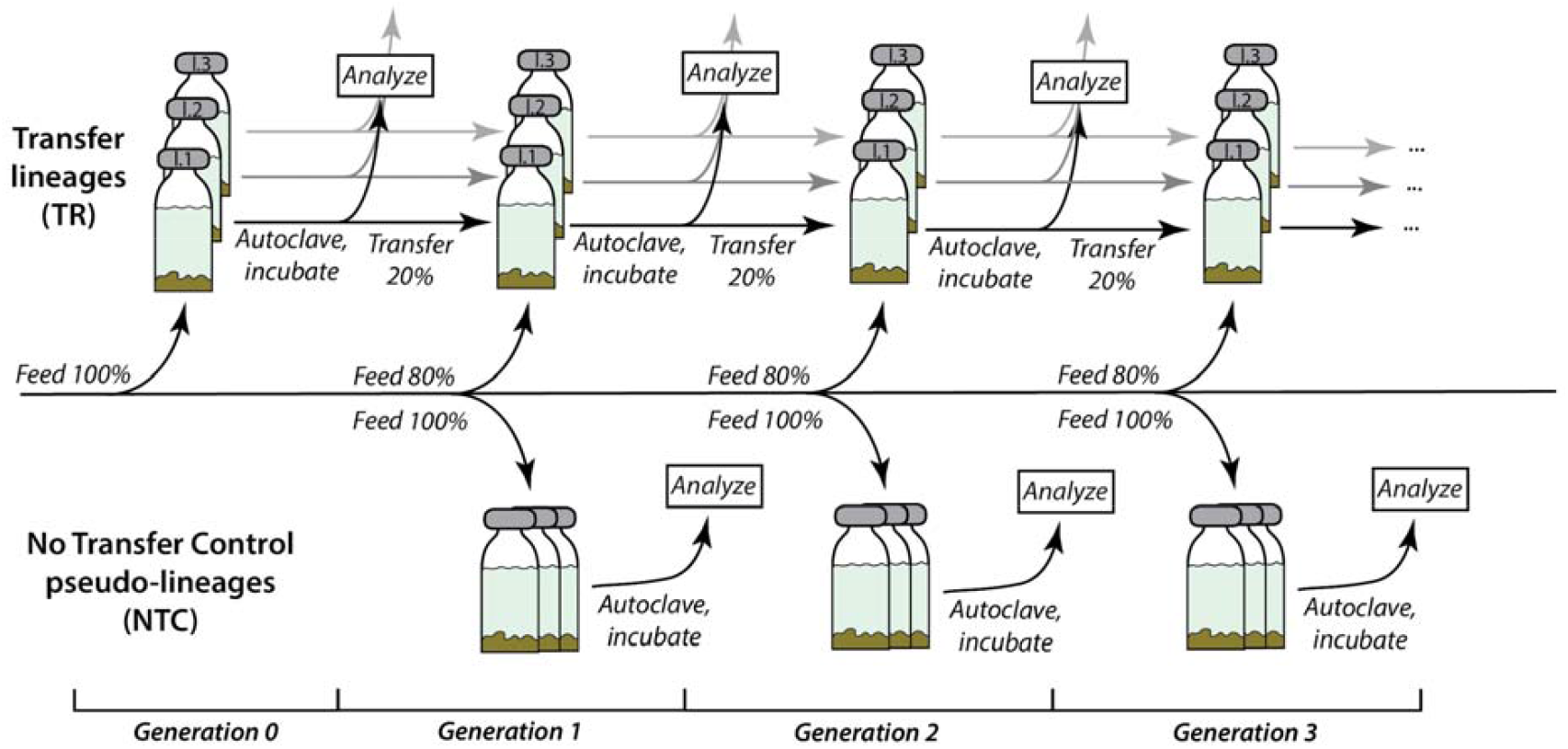
Scheme of the experimental design.

To help interpret the results and explain the role of pyrite and autoclaving in the combinatorial explosion, we examined changes in chemical composition in FS over a single 24-hour incubation period. FS solution was autoclaved and incubated or simply incubated with and without pyrite, with samples taken every 2 hours. In addition to FS, we conducted a similar procedure for samples that lacked subsets of FS components.

Liquid-chromatography/mass-spectrometry (LCMS). Immediately prior to chemical analysis, vial contents were thawed and transferred to 96-well 0.2 mm filter plates (Neta Scientific, PALL-8019) and vacuum filtered, to remove pyrite particles. We used an untargeted metabolomics approach with reverse phase Ultra-Performance Liquid Chromatography coupled with tandem mass-spectrometry (UPLC-MS/MS) with a Thermo Vanquish UPLC (Thermo Fisher Scientific, Waltham, USA) with a C18 column (Agilent, Santa Clara, USA). Samples (5 μL) were eluted in a linear gradient mixture from 0.1% v/v formic acid in water (47146-M6, Thermo Fisher Scientific, Waltham, USA) to 0.1⍰% v/v formic acid in acetonitrile (47138-K7, Thermo Fisher Scientific, Waltham, USA), over 14 min. To reduce the risk that order effects could yield artifactual differences between TR and NTC treatments, samples were run with the first TR and NTC replicates preceding the second replicate of each, and so on.

The UPLC was coupled to a Thermo Q-Exactive Plus Orbitrap MS (Thermo Fisher Scientific, Waltham, USA). Full MS-SIM spectra for each of the replicates were collected for 10⍰minutes in positive mode over a scan range of 50–750 m/z, with resolution set to 70000. Fragmentation data were collected only for pooled samples for each treatment category with full MS followed by data-driven MS2 analysis (dynamic exclusion of 4s, intensity threshold 1.0E5, resolution 17500, isolation window 1 m/z, stepped collision energies of 20, 50 and 100 eV).

Data structure. Data in .raw format was processed in Compound Discoverer™ (Thermo Fisher Scientific, Waltham, USA). We used a default untargeted metabolomics workflow with 16 added databases (Across Organics, Alfa Aesar, Alfa Chemistry, BioCyc, Cambridge Structural Database, CAS Common Chemistry, ChemBank, DrugBank, FDA, Human Metabolome Database, KEGG, MassBank, Merck Millipore, MeSH, NIST, NPAtlas) with pooled samples set to ‘identification only’. Every sample from each experiment was processed in the same Compound Discoverer analysis. Lists of identified compounds, their relative abundances and SMILES IDs were extracted for downstream analyses. Additionally, for samples used to evaluate the role of autoclaving and pyrite in generating chemical complexity, we ran samples individually in Compound Discoverer^™^ in order to obtain comparable counts of the number of MS features. Lists of identified compounds in the combined Compound Discoverer^™^ analysis, their relative abundances and SMILES IDs were extracted for downstream analyses.

All features without assigned names or formulas were removed from the data due to uncertainty about their identity and consequent difficulty in ensuring comparability between samples. Areas of MS features were summed if they had the same name or, if names were absent, the same formula. The resulting dataframes are lists of areas of identified compounds in each sample and their assigned names/formulas.

To evaluate whether the TR and NTC samples in a generation are different, and whether the degree of difference changes over generation, we calculated Canberra (**Eq. 1**) and Bray-Curtis dissimilarities (**Eq. 2**) between each pair of samples using the vegan package in R (Oksanen et al., 2007). These distance metrics are frequently used to quantify dissimilarity between ecological communities (Ricotta & Podani, 2017). Since the Canberra distance sums ratios rather than areas, it tends to weight each species more or less equally, regardless of its absolute abundance, unlike Bray-Curtis which assigns higher weight to more abundant species.

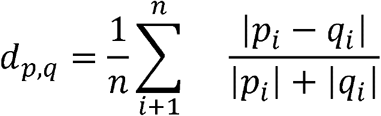

**Equation 1**. Formula for Canberra distance: *n* – number of compounds; *i* – individual compound; *p* – area of compound *i* in one sample; *q* – area of a compound *i* in a different sample.

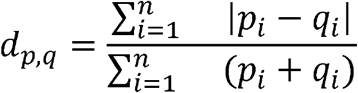

**Equation 2**. Formula for Bray-Curtis distance: *n* – number of compounds; *i* – individual compound; *p* – area of compound *i* in one individual; *q* – area of a compound i in a different individual.

To evaluate whether TR and NTCs differ, we compared the pairwise distances between samples within groups (TRs and NTCs) with the between group distance (NTC-TR). We conducted a permutational analysis of variance (PERMANOVA) in R. We calculated the mean sum of square Canberra and Bray-Curtis distances and obtained standard deviations based on 500 bootstrap resamplings of features.

Heritability calculations. In evolutionary biology heritability is typically estimated as the slope of a linear regression of trait values that have been measured in a set of parents and their corresponding offspring. Since we have 20 parent/offspring pairs in each generation, we have the capacity to calculate an analog of heritability for these data.

The first method we used is similar to the biological method and involves calculating the Pearson correlation of the area of a compound in parents and offspring in adjacent generations (e.g., generations 1 and 2, 2 and 3, etc.). We then compared the regression slope, r, over all compounds to see if the null hypothesis of r=0 can be rejected using a t-test. We also calculated the regression-based correlation for two summary statistics, namely the total number of significantly enriched or depleted compounds relative to the set of NTCs from that generation.

To supplement the biological approach, we also developed a multivariate measure of heritability, the Distance-base Heritability Index (DHI; **Eq. 3**). This index divides the distance between parent-offspring pairs in adjacent generations with the mean distance between non-parent-offspring pairs. Observing DHI values that are lower for TRs than NTCs would show that samples from the same lineage are more similar to each other than would otherwise be expected, thus supporting heritability. Our DHI calculations used the Canberra distance, which weights all compounds equally, regardless of mean concentration.

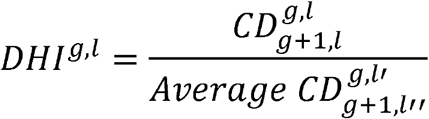

Equation 3. Formula for the distance-based heritability index, *DHI*, as calculated for a parental generation (*g*) relative to its offspring generation (*g* + 1). The average Canberra distance (*CD*) is calculated between each vial in generation g and the corresponding vial from the same lineage (*l*) in generation *g* + 1. This is then divided by the average distance calculated between each parent vial and the offspring vials from all other (non-offspring) lineages (*l*’, *l*’’). This metric is calculated separately for lineages of TRs and pseudo-lineages of NTCs.

Finally, we applied a heredity metric developed by Guttenberg et al. (2015) for use in studies of prebiotic chemistry. We used their Python script (https://github.com/ModelingOriginsofLife/Heredity/tree/master) to analyze our data and estimate the number of heritable states possible for the TRs, as compared to the NTCs. This method conducts principal component analysis on an input set of properties for systems that are inferred to have been exposed to different selective environments. The number of eigenvalues above a computed threshold is taken as an estimate of the number of potentially heritable states. This methodology can distinguish between intrinsic variation due to stochastic processes and extrinsic variation due to selection based on the assumption that intrinsic variation should be unimodal while extrinsic variation could be multimodal (Guttenberg et al., 2015). We assumed that each generation experienced a somewhat different selective pressure due to the differences in the amount of generation 0 material transferred into each vial. Therefore, we estimated the number of heritable states per experimental lineage (N_S_) with the prediction of lower N_S_ in NTCs, which lack true intergenerational transmission. Prior to this calculation, we normalized our dataset by subtracting the mean and dividing by the standard deviation for each compound, as is a common practice for principal component analysis (Guttenberg et al., 2015). The parameter alpha, which sets the sensitivity of the algorithm, was set to 0.1.

Network expansion. To explore the chemical pathways that could have generated the compounds detected with LCMS from the initial FS we inferred a subset of the chemical reactions that are likely occurring (**Fig. 2**) using a new rule-based chemical reaction generation tool, Rule-It (Cuevas-Zuviria & Sokolskyi, 2024). Here, each round of network expansion uses the compounds generated by all previous iterations and the initial food compounds to look for new reactions that are allowable under a defined ruleset. To control the resulting combinatorial explosion, the expansion algorithm in Rule-It allows for arbitrary or data-guided pruning, and can be further constrained by molecular mass of the products (Cuevas-Zuviria & Sokolskyi, 2024; https://github.com/brunocuevas/ruleit).

**Figure 2.**
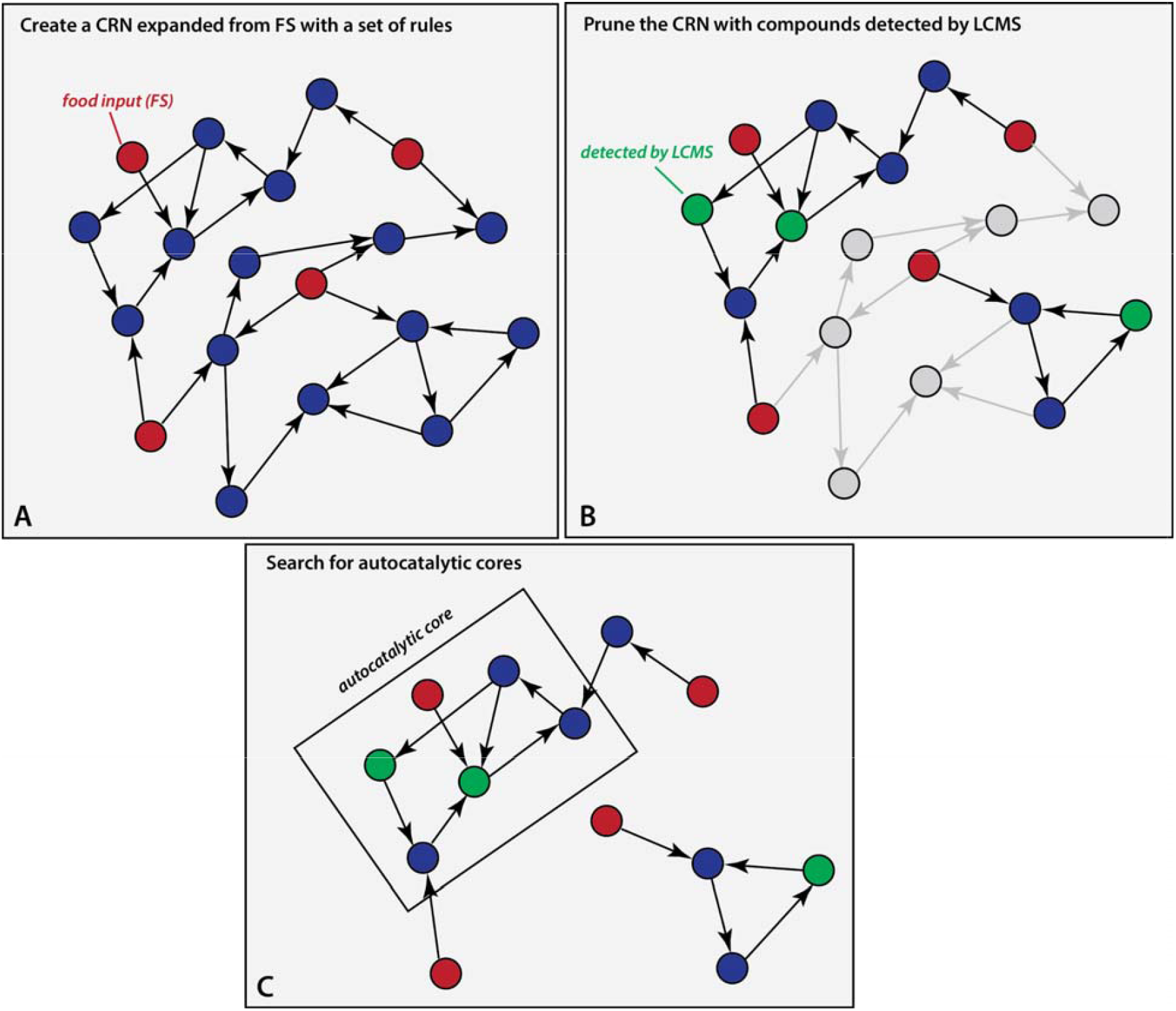
Scheme of the computational pipeline for detecting autocatalytic cores (ACs): first, we expand the network from the initial compound set (FS plus water, sulfate, and hydrogen sulfide), shown in red, using our reaction ruleset (**A**); then we prune the resulting network keeping only reactions and compounds (in blue) that lead from initial compound, to compounds detected with LCMS (**B**); then we search for ACs in the pruned network (**C**).

Our ruleset included 16 reversible reaction types that could plausibly occur in our samples, each written in SMARTS format (**Fig. C1, Tables C1-2**). This ruleset only included reactions with oxygen, nitrogen, sulfur and phosphorus-containing functional groups. To simplify analysis and reduce the total number of allowed reactions, we did not allow any ring-forming reactions, although it is almost certain that some such reactions do occur in these conditions. For computational simplicity, we also did not include rules that would generate the well-known cyanide reaction network (Patel et al., 2015). For the initial set of compounds, we include the six FS solutes plus water, sulfate, and hydrogen sulfide, the latter two being plausible products of pyrite decomposition. Using a 40000 reaction limit and molecular weight cap of 500, we ran four iterations of network expansion (**Fig. 2, A**).

To reduce the size of networks to a manageable size while focusing on potentially relevant compounds, we pruned networks guided by experimental evidence. We identified all compounds in the network that were detected by LCMS and pruned the network to include these species and all reactions on the shortest reaction path to these compounds from the initial set of compounds. All of the input files are available in Supplementary data.

Detection of autocatalysis. To detect autocatalytic cores (ACs), or minimal autocatalytic motifs, in the pruned FS-driven CRN we utilized the program autocatatalyticsubnetworks (Gagrani et al., 2024), which is downloadable from Github (https://github.com/vblancoOR/autocatatalyticsubnetworks). Using that software, we converted the pruned networks into stoichiometric matrices and then analyzed these to detect stoichiometrically autocatalytic motifs (**Fig. 2, C**). To reduce computation time, we removed rows representing water, carbon dioxide, ammonia and hydrogen sulfide from the input matrix, thereby preventing us from detecting autocatalytic cores with these compounds as member species. We did this because ACs with these ubiquitous compounds are poor candidates for explaining heritability.

## Results

### Pyrite and autoclaving promote chemical diversity

To test for the existence of heritability, it was necessary that the experimental system generated sufficient chemical complexity that different lineages could, in principle, diverge from one another. Based on individual Compound Discoverer^™^ runs we evaluated the average number of features above an arbitrary area threshold of 10^6^ for different treatments (**Supp. Table B1**). The largest factor seemed to be the presence/absence of pyrite since more than 500 features were detected in samples with FS and pyrite, compared to 166 without. The massive increase in the number of compounds seen after incubation with pyrite with or without autoclaving can plausibly be explained by the chemistry of the food as summarized in **Appendix A**. However, almost 500 features were also detected in samples with water + pyrite, which might suggest that the effect of pyrite is partly due to contamination. While we cannot rule this out, it is noteworthy that, almost all compounds found in water + pyrite are undetected or at low abundance in FS + pyrite (**Fig. 3**). As result, it is possible many represent analytical artifacts rather than contamination from pyrite. Sodium and potassium salt clusters and other metal ions are a common electrospray ionization (ESI) artifacts (McMillan et al., 2016), and likely correspond to many detected features, since FS includes two sodium salts (bicarbonate and trimetaphosphate). Differences in pyrite oxidation between water and FS could also affect feature counts as Fe adducts are known to be another ESI artifact (Bonner & Hopfgartner, 2022). Hence it is unclear whether absolute feature counts accurately represent chemical diversity of our samples.

**Figure 3.**
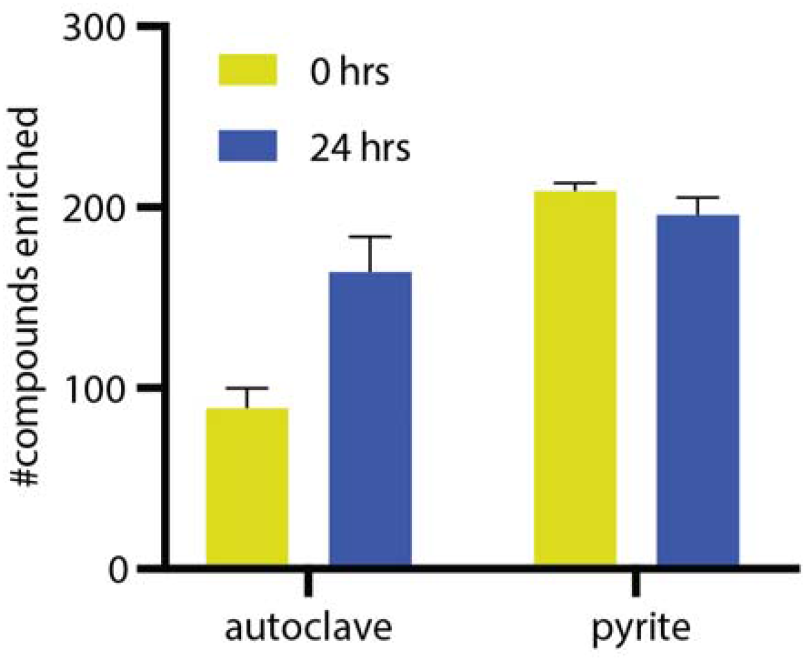
Number of detected compounds with area >10^6^ significantly enriched relative to respective controls (>2*SD difference: compared to unautoclaved samples without pyrite for autoclave effect; compared to unautoclaved samples without pyrite and to water with pyrite for pyrite effect) in FS due to autoclaving or pyrite. Error bars are SEM.

Combined analysis in CompoundDiscoverer™ makes it possible to directly compare the mass spectral features between samples, at the cost that all compounds that are detected in any sample are assigned an imputed area in all other samples, whether or not they were detected. To get around this problem and quantitatively assess how pyrite and autoclaving affect compound abundances, we counted up the number of compounds that have significantly elevated areas in one treatment relative to another (**Fig. 3**). The effect of autoclaving was assessed by comparing FS without pyrite to autoclaved FS over the course of a 24 hour incubation. The effect of pyrite was determined by comparing unautoclaved FS with pyrite to unautoclaved FS without pyrite. These analyses revealed that an “explosion” of chemical diversity occurred both due to either autoclaving or pyrite addition, with pyrite having a greater and more immediate effect.

Transfer samples diverge from controls. To quantify the divergence between transfer lineages and control lineage, we conducted PERMANOVA analysis on Canberra and Bray-Curtis distances. For Canberra distances NTC-TR distances are greater than either NTC-NTC or TR-TR distances, except for generation 6 when TR-TR distances are higher. This difference was significant (p<0.05) in every generation except generation 5 (**Fig. 4, A**). Bray-Curtis distances also tend to be more similar within than between groups, but NTC-TR and NTC-NTC are significantly different only in generations 3, 4 and 6 (**Fig. 4, B**). Taken together, the tendency for greater NTC-TR distances than within-group distances shows that the presence of material transferred from prior generations significantly affects the overall composition of vials.

**Fig. 4.**
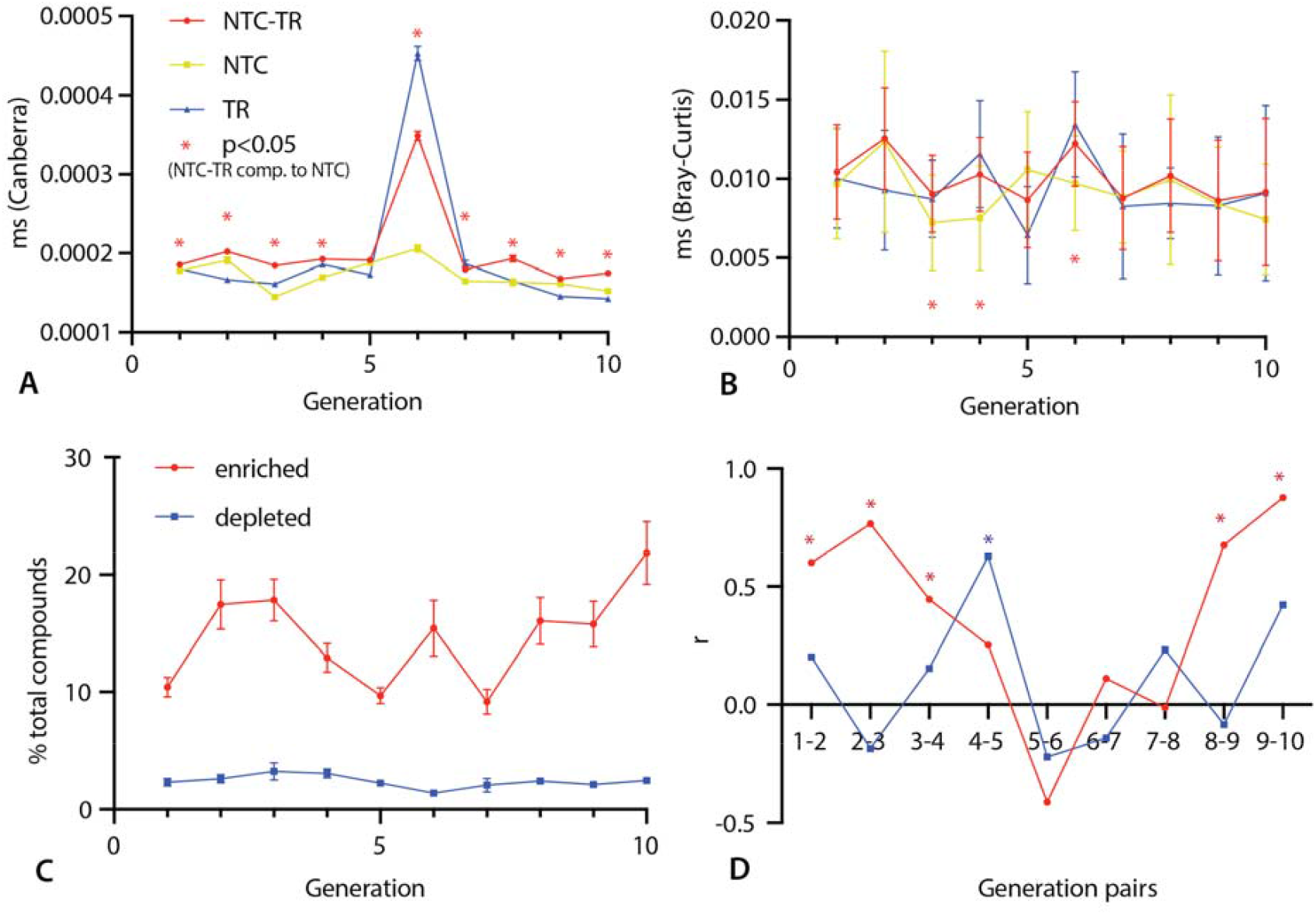
Dynamic changes over the course of the experiment. **A** – mean sum of squares (ms) of inter-group Canberra distance, plotted are TR-NTC, NTC-NTC and TR-TR group distances; **B** – mean of squares (ms) of inter-group Bray-Curtis distance, plotted are TR-NTC, NTC-NTC and TR-TR group distances; **C** – change in the number of compounds significantly (difference greater than 2* SD_NTC_) enriched or depleted in TRs compared to NTCs every generation; **D** – Pearson correlation of the number of enriched and depleted compounds between adjacent generations in the TR treatment (significant correlation, p<0.05, is marked by asterisks). Error bars are SD in **A-B** and SEM in **C**.

In addition to distance metrics, we also calculated the number of compounds that are significantly (>2*SD_NTC_ difference) enriched and depleted in TRs compared to NTCs within a generation (**Fig. 4, C**). Since TRs carry forward products from one generation to the next, we predicted that there would often be compounds that are only seen above the threshold of detection in TRs. Indeed, about ten times as many compounds are enriched than depleted when comparing TRs and NTCs. While the number of depleted compounds remains similar across all 10 generations, the fraction of compounds that are significantly enriched in TRs varies markedly across generations (9-22%).

We calculated the Pearson correlation coefficients between the number of enriched and depleted compounds in parent-offspring pairs (**Fig. 4, D**). While the slope of the regression of depleted compounds fluctuates, there is generally a positive correlation for enriched compounds, meaning that parent vials with a higher or lower count of enriched compounds tend to give rise to offspring vials that also have a higher or lower number, respectively. This parent-offspring correlation has a significant slope (p<0.05), suggesting heritability, for five generation pairs, the first three and the last two (**Fig. 4, D**).

### Heritability of chemical composition

More individual compounds have significantly positive parent-offspring correlations than expected by chance, but a similar fraction was seen in both TRs and NTCs despite the fact that NTCs have no true “parent-offspring” relationships. Similarly, although average regression slopes are significantly greater than 0 in many generations, these were found in both TRs and NTCs with no clear pattern (**Fig. B1**). The most likely explanation of these results is not heritability but the order in which samples were analyzed in LC-MS. Lower-numbered replicates of both NTCs and TRs tended to be loaded earlier in an LC-MS run than high-numbered replicates, which can yield a false signal of heritability when column properties or MS sensitivity changes over time.

In order to obtain a single metric that summarizes all data, we developed the DHI statistic, which measures whether parent-offspring pairs are more similar than unrelated vials from their respective generations. When calculating DHI using the Canberra distance matrix for the complete, filtered dataset we observed no difference between TR and NTC values, which are all in 90-100% range, except for generation 1, where the DHI of TRs is significantly (p<0.05) greater than in NTCs (**Fig. 5, A**).

**Fig. 5.**
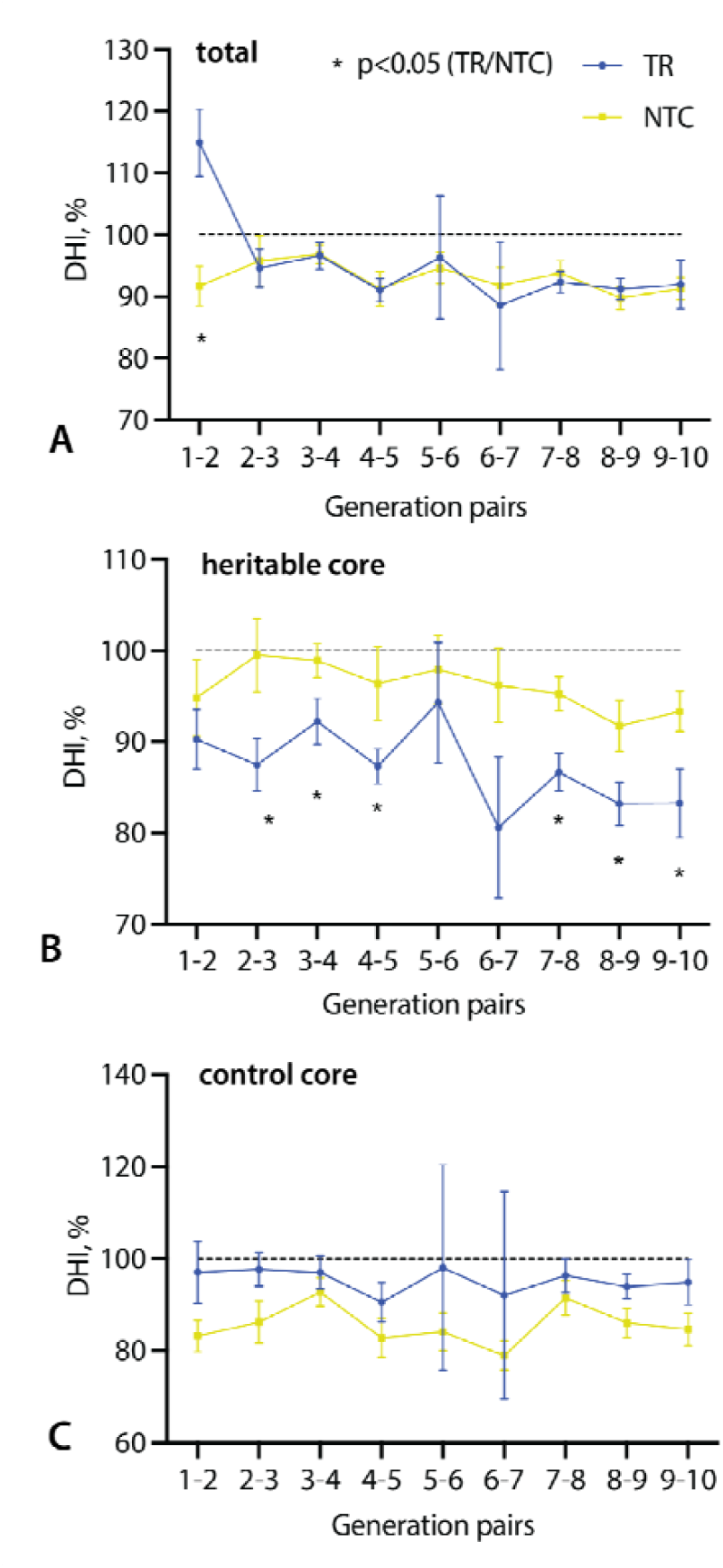
Changes in heritability as measured by the distance-based heritability index (DHI) using Canberra distance matrix. **A** – change in DHI over the course of the experiment for the whole compound dataset; **B** – change in DHI for the dataset comprising the “heritable core” – compounds that have r>0.5 in 3 or more pairs of generations of TRs, and 1 or less generations of NTCs; **C** – change in DHI for the control dataset – compounds that have r>0.5 in 1 or less pairs of generations of TRs, and 3 or more generations of NTCs. Error bars are SEM.

Given the possibility that a strong order-of-analysis effect might be swamping true signals of heritability we recalculated DHI based only on compounds that are the strong candidates for being heritable in TRs but not NTCs and, as a control, those that are candidates for showing higher “heritability” in NTCs. To be a candidate in either direction, a compound showed Pearson correlation coefficients greater than 0.5 in 3 or more pairs of generations for one of the data sets (TR or NTC), but for 1 or fewer pairs of generations for the other data set. Recalculating DHI for the heritable candidate sets yielded different results for TR and NTC data (**Fig. 5, B-C**). In both cases the DHI for the candidates is reduced as expected if there were heritability. This could be due, in part, to selecting those compounds that had a pattern of apparent heritability. However, the DHI decline was much greater when the compounds were selected based on apparent heritability in the TRs (**Fig. 5B**) than for the NTCs (**Fig. 5C**). Statistical tests never detected a significant difference in DHI between NTCs and TRs for the NTC candidates, but the TR DHI is significantly lower than those for NTCs in 6 out of 9 generation pairs for the TR heritable core (**Fig. 5, B-C**).

We also estimated the number of possible heritable states for the full dataset and the two candidate subsets, using the method of Guttenberg et al. (2015). For the total dataset we find that TR lineages have more possible heritable states with a close to significant p-value of 0.06 (**Fig. 6**). For the TR candidates the difference is even more significant (p = 0.02), while for the NTC candidate no significant difference was detected (p=0.37).

**Fig. 6.**
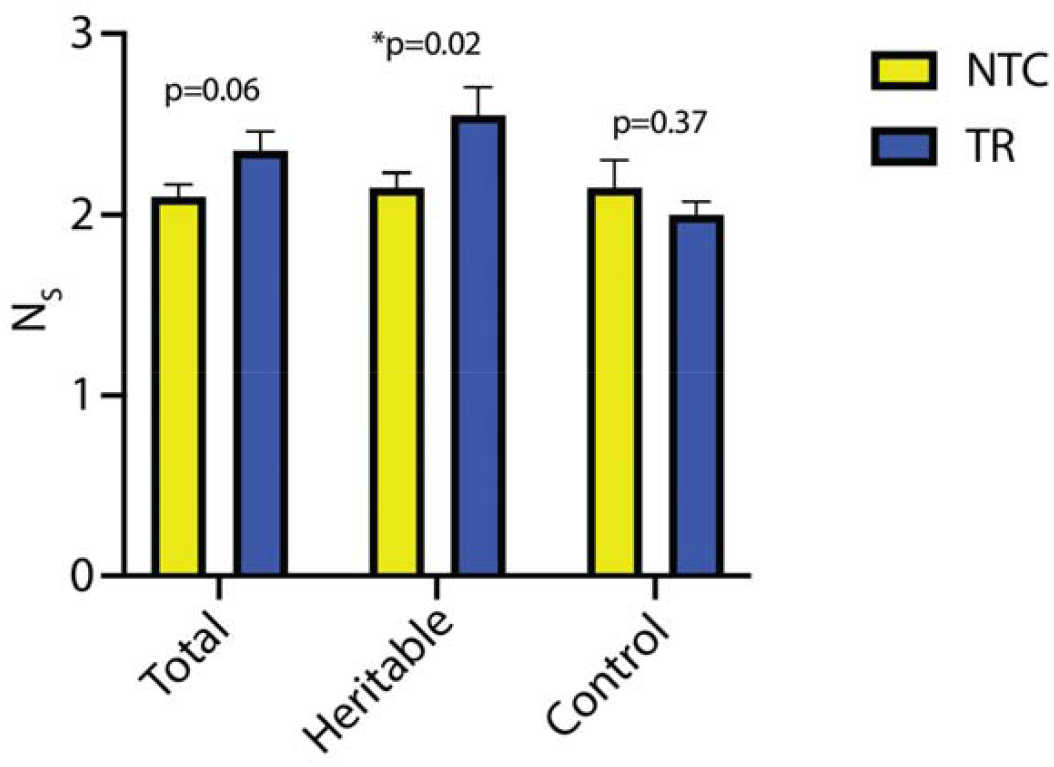
Number of heritable states (N_S_), for the total dataset, heritable core (only compounds with r>0.05 for >3 pairs of generations of TRs and <1 of NTCs) and control core (only compounds with r>0.05 for >3 pairs of generations of NTCs and <1 of TRs). Error bars are SEM.

### CRN reconstruction and analysis

Using the FS as the input and the ruleset shown in Table C2, one round of expansion yielded a CRN of 22 reactions and 28 compounds (**Fig. C2**). This increased to 332 reactions and 282 compounds after two rounds, 7869 reactions and 6719 compounds after three, and 40395 reactions and 33873 compounds after 4. In the latter CRN, 47 compounds could be matched to features that were detected and identified by LCMS. Of the 6 (1.5%) or the 412 compounds that were candidates for heritability based on the NTCs and 3 (1.9%) of the 160 compounds in the TR candidates were included in the network. Such low numbers are expected, as our ruleset likely represents a small fraction of possible FS-driven reactions. The numbers of reactions for each rule are summarized in supplementary **Fig. C3**.

In the resulting CRNs pruned on either all 47 detected compounds or the two candidate sets, we detected multiple autocatalytic cores (ACs), each including from 10 to 34 reactions. There were 419 ACs for the network pruned with all LCMS hits, 1 for the NTC heritable candidates, and 14 for the TR heritable candidates. Many of the ACs included nitrogen- or sulfur-containing compounds, some included only carbon, oxygen, and hydrogen. An example of the latter is shown in **Fig. 7**. We were unable to detect any ACs in the pruned network whose food consisted only of compounds in the FS, meaning that at least one food has to be produced by other reactions that are not part of the AC. However, for many of the ACs the food compounds are relatively simple, e.g., formic acid, small imines, or methanol. Additionally, because some key reactions are removed during network pruning, there are likely additional pathways for generating some food species. For example, R07 in **Fig. 7** was not present in the pruned network - and thus the AC had species (1) as a food and species (9) as a waste species. However, R07 was present in the full network resulting from network expansion.

**Figure 7.**
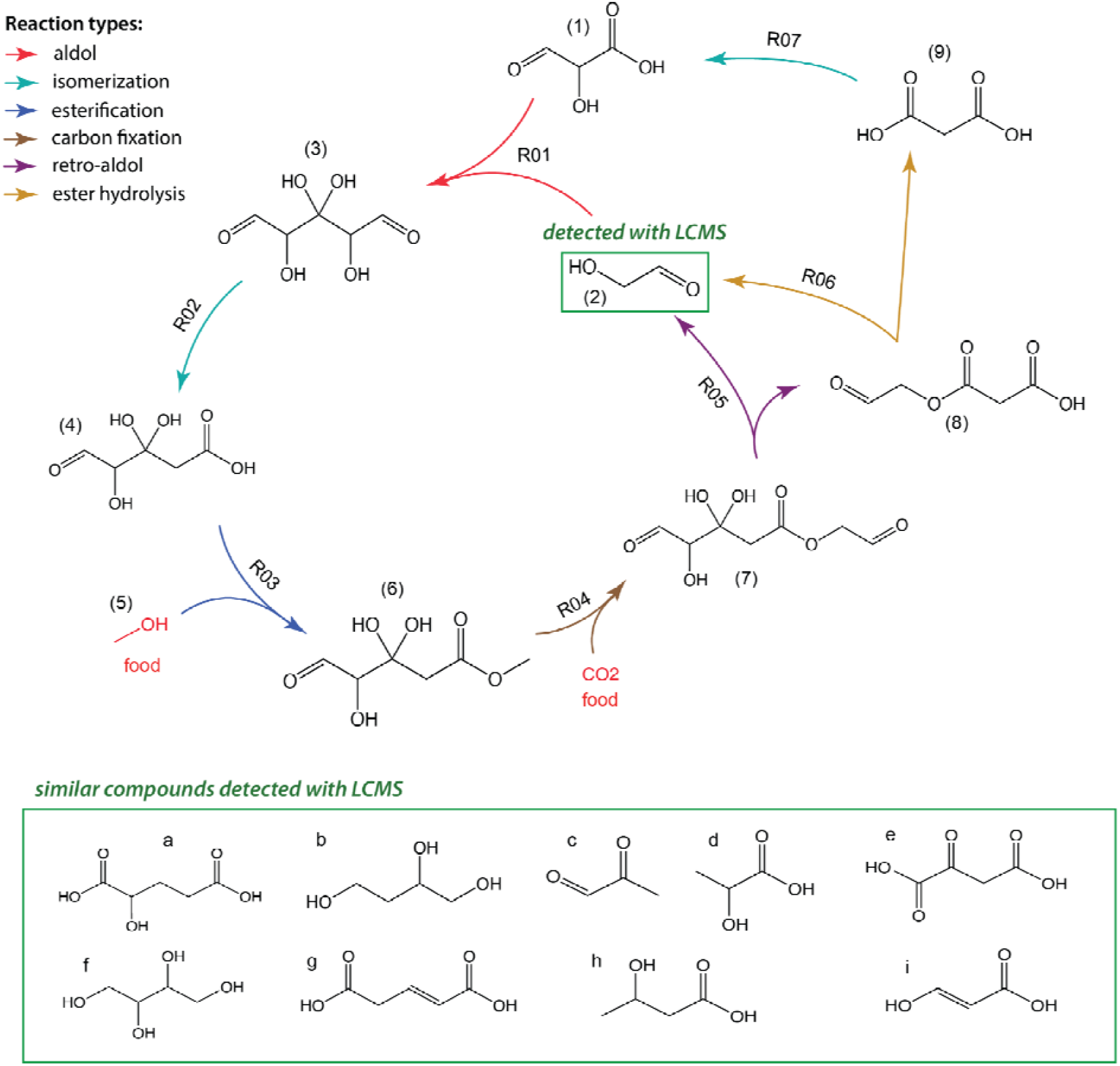
An example AC detected in the CRN pruned with heritable candidates from the TR.

For none of the detected ACs were all of the member species detected and identified by LCMS. This could of course be due to analytical constraints, but could also be the result of certain member species having low equilibrium concentrations, for example due to them being unstable intermediates. However, in many cases chemically similar species were detected. For the AC in **Fig. 7** only glycolaldehyde (2) was detected but there are multiple alcohols, aldehydes and carboxylic acids in the LCMS data similar to other AC member species. For example, compound d (lactic acid) only differs from compound 1 (2-hydroxy-3-oxopropanoic acid) by one carbonyl group.

## Discussion

### Autoclaving, pyrite and transfers significantly impact chemical diversity

Both autoclaving and pyrite result in the appearance of many more compounds in FS (**Fig. 3**). Pyrite had the greatest effect, presumably because of the ability of pyrite and other metal-rich minerals to catalyze prebiotically-relevant reactions, including abiotic carbon fixation, incorporation of nitrogen into organic compounds, and the formation of larger organics (de Graaf et al, 2023; Asche et al., 2024). Carbon fixation with CO_2_, or similar reactions starting with 1 and 2 carbon organic molecules, seem particularly relevant and necessary to explain the many larger organic compounds we detected after incubation. As outlined in Appendix A, we can draw on changes in vial composition over the course of a 24-hour incubation to speculate as to a few chemical reactions that might be occurring during incubation with pyrite.

In order to have any possibility of revealing heritability, it is first essential that the transfer process affects the chemical composition of TR vials relative to NTC vials, which receive no inputs from the prior generation. Indeed, we find clear evidence of systematic differences between TR and NTC vials in almost all generations, as shown by greater average TR-NTC Canberra distances than either TR-TR or NTC-NTC (**Fig. 4, A**). The fact that this effect is less clear using Bray-Curtis distances (**Fig. 4, B**) likely reflects the fact that this distance measure tends to be dominated by a few high-abundance compounds.

### Heritability can be detected in small-molecule chemical systems

In this study we used multiple parallel parent-offspring transfer lineages to detect heritable variation emerging from prebiotically plausible reactions. Here we understand heritability, analogously to biology, as the fraction of the variation between individuals in a population that is inherited from parents to offspring (Visscher et al., 2008). In this context, the 20 TR vials in adjacent generations comprise the population of parents and offspring and the phenotypes are areas of different mass-spectral features.

To be relevant to long term evolution, heritable differences among vials would need to be the result of autocatalytic cycles that are activated in some but not all vials. However, any given autocatalytic cycle may have only a small number of detectable members meaning that only a tiny fraction of compounds will show heritability, with the remaining variation being due to experimental noise. Thus, the fact that we did not find many compounds in TR vials that had high parent-offspring correlations is insufficient to reject the hypothesis of heritability.

Three analyses supported the idea that heritability is present in this system. First, a system-level, emergent trait, the number of enriched compounds, shows a pattern consistent with heritability (**Fig. 4, D**). Second, we found DHI to be significantly higher for the TR heritable candidates in most generations, whereas no such pattern is seen using the NTC candidates (**Fig. 5**). Third, the number of heritable states in TRs tends to be significantly greater than in NTCs (**Fig. 6**). The latter is more a metric of evolvability than heritability, and may be sensitive to the larger number of enriched compounds in TRs than NTCs, but seems to confirm the general pattern that transfer lineages have the potential to diverge from one another over time. Collectively, these analyses provide the first direct evidence that chemical systems driven from equilibrium by prebiotically plausible chemical food sets can vary and pass on those variations into the future. This is important because the generation of heritable variation is a minimal requirement for evolution by natural selection.

To evaluate the significance of this result, it is worth considering whether these patterns could arise by trivial mechanisms, of which the most obvious is contaminants being introduced into a subset of parent vials that are then seen at elevated concentrations in offspring vials simply because the latter receive, as input, 20% of the volume from their parent vial. If such contaminants did not react, we would expect them to occur at five-fold lower concentration in offspring than parents. A positive correlation between parents and offspring could be seen, nonetheless, but only if the magnitude of the contamination received by parents is five times greater than the threshold of detection by LC-MS. We believe that such high-concentration contaminants are likely rare, so we take the evidence of heritability as suggestive of autocatalysis.

### Autocatalysis and heritability

Our hypothesis is that variation in parent vials arises from time to time, whether from sub-threshold contamination or stochastic chemical dynamics, and this triggers novel autocatalytic processes that are passed onto offspring vials. Consistent with this, we detected multiple stoichiometric ACs in the partial CRN that we inferred using a limited number of reaction rules, while disallowing cyclization reactions. Moreover, even with these constraints, the number of ACs detected was surely a large underestimate. First, due to the probabilistic nature of the expansion algorithm, some important reactions would have been missed. This is illustrated by R07 in **Fig. 7**, which was present in the unpruned network, but was removed after pruning. Second, Rule-it only considers uncatalyzed unit reactions, disallowing cases of catalysis, where the same compound appears both as a reactant and product. Third, we were forced to limit the network expansion to just four iterations due to computational constraints, whereas the number of autocatalytic motifs tends to scale exponentially with CRN size (Mossel and Steel, 2005; Gagrani et al. 2024). These factors suggest that the number of ACs we detected is a significant undercount.

A promising finding that could be taken as supporting our inference, is that the CRN pruned based on the 3 members of the heritable core set contained 14 ACs, as contrasted with just one in the network pruned based on the NTC candidate set. If heritability is associated with autocatalysis, we expected the TR heritable candidates to include members ACs - while AC members in the NTC candidates must be random. However, we must be cautious about this observation, as it could easily be a stochastic artifact due to the small number of compounds detected in each case.

To support heritability, it is not sufficient that stoichiometric ACs be present, they also would need to be kinetically viable (Steel et al., 2019). For example, a food species for an AC may not be produced at a sufficient rate to allow for exponential growth of member species or side reactions may consume some member species at rates greater than the AC reactions (Orgel, 2008). Thus, it would be desirable to obtain data on plausible rate constants of different reactions or, better still, to experimentally validate inferred ACs.

Even if kinetically viable, an AC could only support heritability if it differed in flux between lineages, the extreme being the case where it is active only in a subset of vials that were seeded by slow reactions or occasional contamination (Peng et al., 2022). Whether or not this can be experimentally shown, it highlights the potential importance of rare events and chemical diversity in prebiotic evolution, which contrasts with the focus on high-yield reactions that lead to particular target molecules, which has dominated much prior research in prebiotic chemistry.

This work provides the first statistical evidence for the spontaneous emergence of heritable variation in prebiotically plausible chemical mixtures. Additionally, we developed a novel set of methods for quantifying heritability and connecting it with the underlying chemical reaction network. This represents a significant advance towards the goal of understanding how systems capable of adaptive evolution could have bootstrapped themselves into existence on the early Earth. Moreover, while there are areas where the protocol could be improved, most obviously by randomizing LC-MS analysis order to remove order-of-analysis effects, our experimental approach and the metrics developed may serve as a blueprint for further investigations into the emergence of heritability and evolvability in the absence of genetic polymers.

## Acknowledgments

The authors would like to thank Zoe Todd, John Yin, Betül Kaçar, Daniel Amador-Noguez and Annie Bauer for feedback on this project. We would like to thank Chris Thomas, Julia Duncan and Eric Armstrong for valuable guidance on LCMS operation and data analysis. We would like to thank Praful Gagrani for clarifications on mathematical concepts and Lena Vincent for help with experimental design. We thank Cécile Ané from the Statistical Consulting group at the University of Wisconsin – Madison for assistance in the analysis of the experiment.

## Data availability

Data and code used in this study are available free of charge at DataDryad https://doi.org/10.5061/dryad.hhmgqnkrt.

