## Appendix for "From messy chemistry to ecology: autocatalysis and heritability in prebiotically plausible chemical systems"

**Appendix A. Characterization of the possible chemistry of the food set**

This appendix shows results from preliminary experiments with the food set (FS) solution used in the experiments of this study. FS is composed of 6 soluble compounds (acetic acid, formic acid, methanol, ammonia, sodium bicarbonate and sodium trimetaphosphate) and pyrite in an aqueous solution at pH 7. In addition to long-term transfer experiments, we analyzed chemical composition of FS under different conditions: with and without pyrite; with and without autoclaving; and with partial compositions. Results of these analyses are presented here.

It is important to note that these data is by no means an exhaustive characterization of the FS chemical diversity. With our LCMS analytical techniques derived from untargeted metabolomics some compounds may be more frequently detected than other, biasing the overall results. We also have not utilized any additional analytical techniques on these samples, so the discussion below of possible reactions occurring in our samples is largely hypothetical at this stage.

**Products of pyrite oxidation & production of sulfur-containing compounds.** When in an aqueous environment and in the presence of molecular oxygen, pyrite (FeS_2_) can oxidize to produce various iron and sulfur oxides (Feng et al., 2019). While our experiments were conducted under anaerobic conditions (95% N_2_, 5% H_2_), trace amounts of oxygen were still present (10-20 ppm). For example, dissolution of pyrite in oxygen-containing water can precipitate goethite (Bladh, 1982). High temperature and pressure, as present in the autoclave, can also affect pyrite decomposition, as pyrite can transform to pyrrhotite (Fe_1-x_S) and then to troilite (FeS), releasing sulfur radicals capable of forming H_2_S (Ma et al., 2021). At temperatures greater than 300C, pyrite can be the main source of hydrogen disulfide during in-situ conversion of shales (Kelemen et al., 2012). Experiments involving heating pyrite show H_2_S evolution is minute at ~300C and starts to increase only after ~340C (Ma et al., 2021), and this reaction is thermodynamically possible starting at 275C (Ding & Liu, 2017). However it is still likely H_2_S is produced in our experiments at least in trace amounts due to recursive autoclave cycles and high pressure conditions, which should increase the rate of pyrite decomposition.

Over the course of 24-hour incubation tests and the long-term experiments we detect multiple compounds likely associated with pyrite decomposition: goethite, iron (II) oxide, sulfate anions, thiols, thials, sulfonic acids, sulfides and other organic sulfur-containing compounds. It was shown previously that thiols can form from FeS/FeS_2_ redox system under comparatively mild conditions (Kaschke et al., 1994) and subsequently form thioesters (Weber, 1998). Other sulfur-containing functional groups can be formed in our system either through thiol oxidation or reactions with sulfate anions, e.g., catalyzed by metal oxides (Wallace, 1966). Additionally, in the presence of FeS and H_2_S, the resulting FeS/FeS_2_ redox couple can potentially catalyze production of numerous organic compounds through electrochemical gradients (Wächtershäuser, 1988; Murphy & Strongin, 2009).

In the 24-hour incubation tests, with and without autoclaving we find that autoclaving has an effect on pyrite oxidation (**Fig. A1**). Unautoclaved samples show less iron (II) oxide (**Fig. A1, A**) and more metallic iron (**Fig. A1, B**), implying that autoclaving increases the rate of iron oxidation. Abundance of either compound does not change over the course of 24 hours. However, goethite, a Fe^3+^ pyrite oxidation product, is considerably more abundant in unautoclaved samples, especially at the start of incubation, following which its abundance decreases (**Fig. A1, C**). Concentration of sulfates is independent of autoclaving or incubation time, implying that most sulfate production occurs quickly upon pyrite addition to FS (**Fig. A1, D**). **Fig. A1, E**-**F**, a thiol and a sulfonic acid, are two examples of organic sulfur-containing compounds consistently detected in our samples, abundance of which does not change due to autoclaving. Some other sulfur-containing compounds, such as ferroglycine sulfate, seem to decompose due to autoclaving (**Fig. A2**).

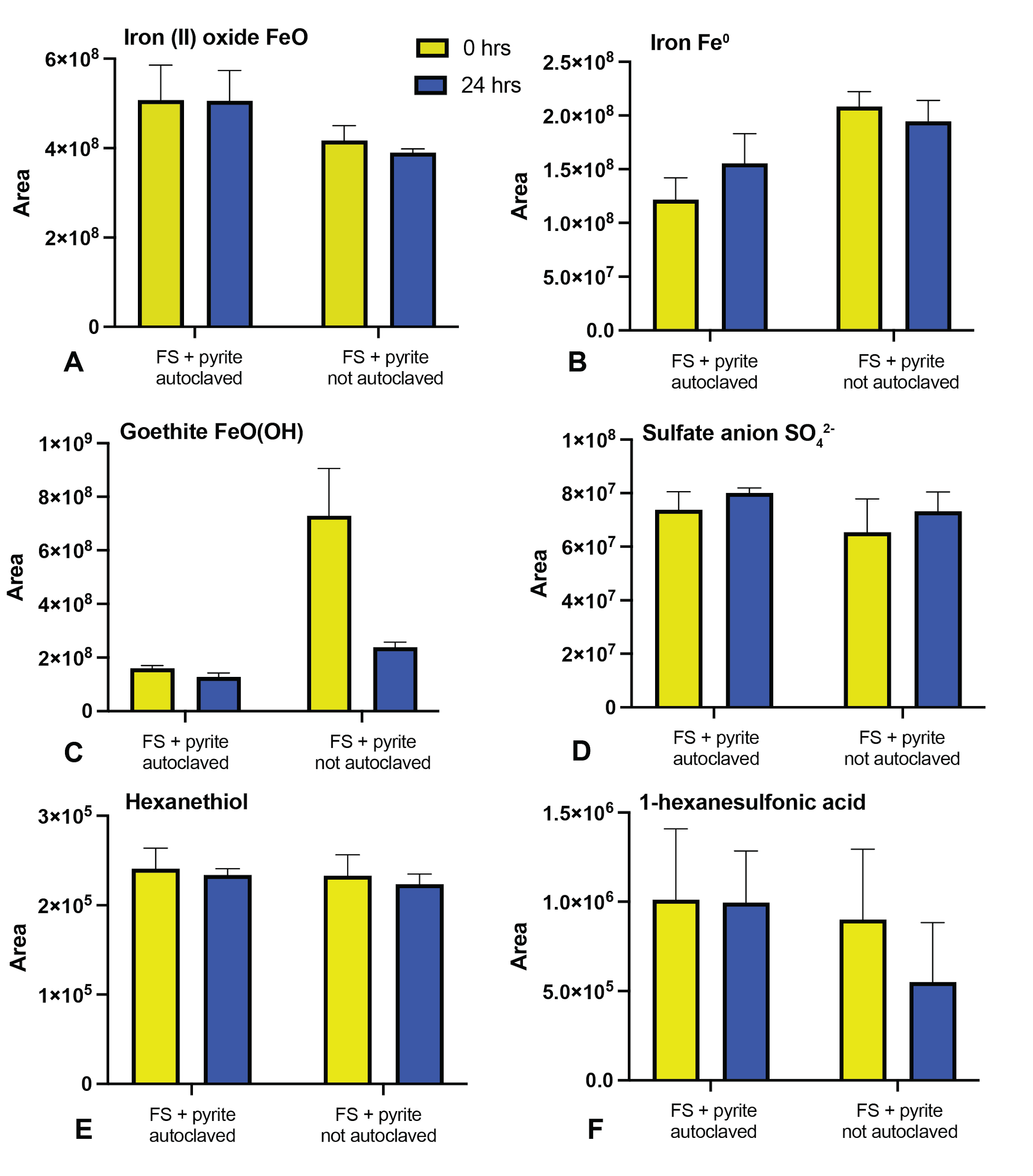

**Fig. A1.** Relative abundance of different iron- or sulfur-containing compounds in the 24-hour incubation test of the FS with pyrite and/or autoclaving. A – iron (II) oxide; B – iron; C – goethite; D – sulfate anion; representative organic compounds: E – hexanethiol; F – 1-hexanesulfonic acid. Error bars are SEM.

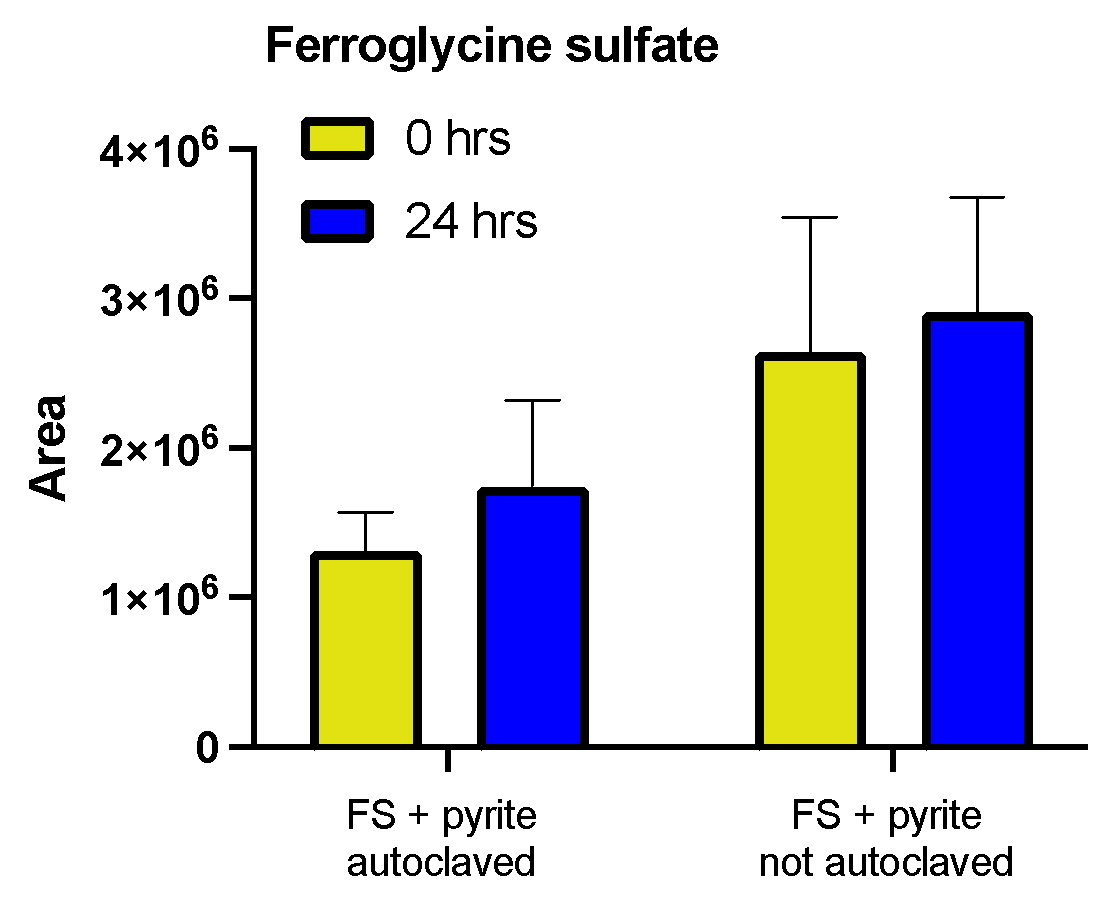

**Fig. A2**. Relative abundance of ferroglycine sulfate changing due to autoclaving and incubation. Error bars are SEM.

Notably, composition of FS has an impact on pyrite decomposition (**Fig. A3**). We observe the greatest abundance of iron oxides and sulfate in samples containing organic FS components (formic, acetic acids & methanol) and ammonia, meanwhile bicarbonate, inorganic FS mixture and FS total have the lowest. This might imply that bicarbonate specifically may have some protective effect, decreasing the rate of pyrite oxidation. However, available literature suggests that carbonate-iron (II) complexes promotes pyrite oxidation to iron (III), which we do not observe with iron (III) species detected in our system (Evangelou et al., 1998; **Fig. A3, B**). An alternative, more likely, explanation is that other FS components increase pyrite oxidation to a greater extent than bicarbonate alone. It is also possible that some pyrite oxidation products are simply not detectable with our methodology.

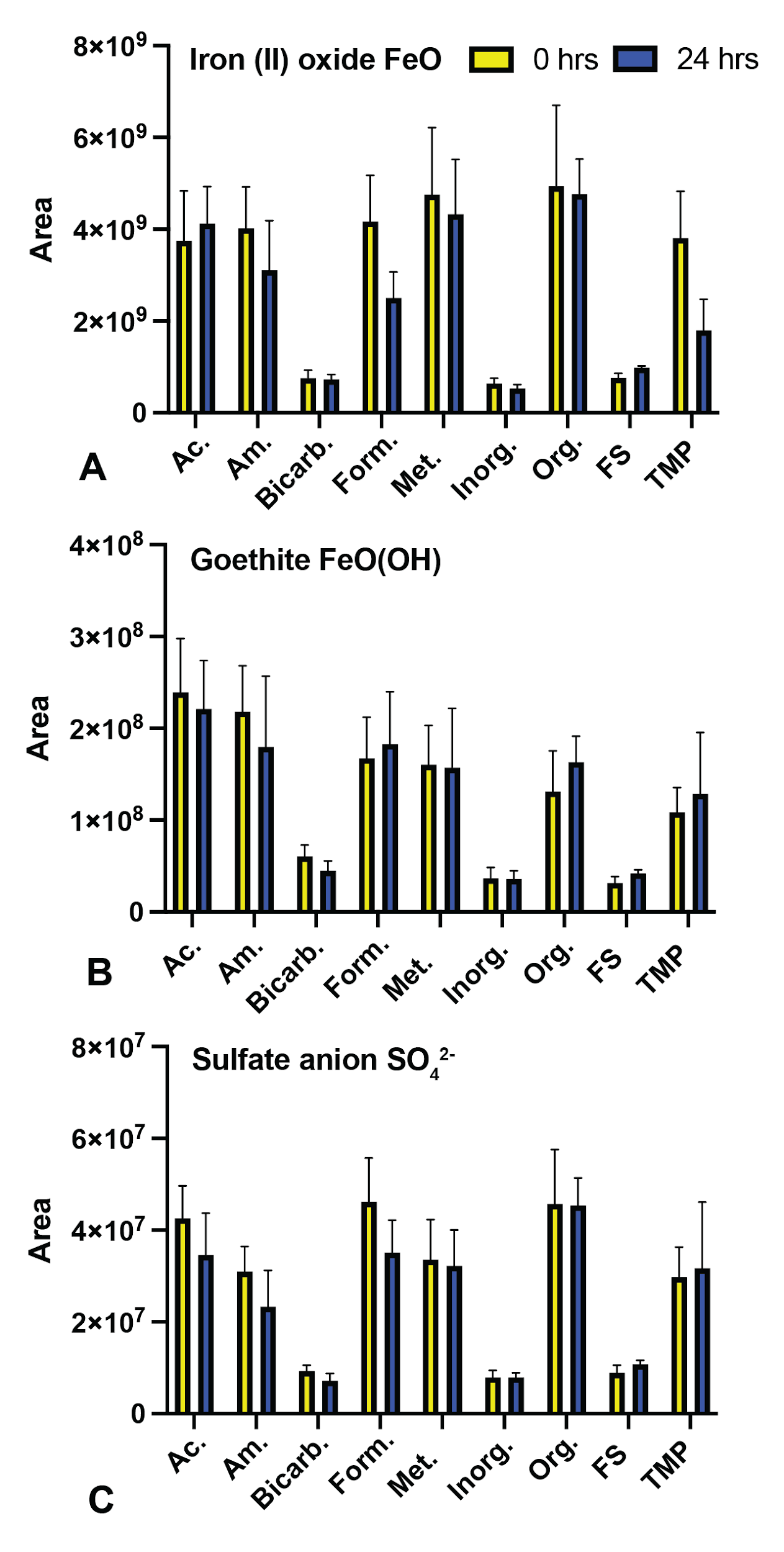

**Fig. A3**. Relative abundance of iron (II) oxide (A), goethite (B) and sulfate (C) in autoclaved samples with pyrite with different composition: Ac. – only acetic acid; Am. – only ammonia; Bicarb. – only sodium bicarbonate; Form. – only formic acid; Met. – only methanol; TMP – only sodium trimetaphopshate; Inorg. – only inorganic FS components (ammonia, bicarbonate, TMP); Org. – only organic FS components (formic, acetic acids and methanol); FS – complete food set. Error bars are SEM.

**Formation of organic molecules.** Another potential effect of pyrite in our system is catalysis of various organic reactions, including carbon fixation (de Graaf et al., 2023). Previous studies have found both high temperature conditions and presence of pyrite surface conducive to producing longer carbon chains (Wachterhauser, 1988; Cody et al., 2004; Michalkova et al., 2011; Camprubi et al., 2017 etc.). Pyrite and other transition metal sulfides are known to promote hydroxycarboxylation – insertion of a carbonyl group at a metal sulfide-bound alkyl group (Cody et al., 2004). Nickel-iron nanoparticles can catalyze formation of formate, acetate and pyruvate from carbon dioxide (Bevazay et al., 2023; Belthle et al., 2023). A variety of sulfur-containing organic compounds was detected in anaerobic reactions of FeS, H_2_S and CO_2_ (Heinen & Lauwers, 1996). Alternative carbon sources were also shown to react with pyrite, for example, methanol and KCN mixtures (Hennett et al., 1992) and pyruvate with ammonia (Novikov & Copley, 2013) yielded amino acids and formic acid to pyruvate conversion (Cody et al., 2000). Autoclaving may contribute to these reactions as well – it was shown to result in production of organic compounds from CO_2_ together with meteoritic and volcanic particles (Peters et al., 2023).

Overall, it is reasonable to expect a high diversity of organic compounds from the FS mixture – which is what we observe. We also find that the number of compounds containing 2 or more carbon atoms (the longest organic compound in FS is acetic acid, at 2 carbons) is significantly greater in FS with pyrite than without and is seemingly independent of autoclaving (**Fig. A4, A**). Additionally, this number is greater in samples without bicarbonate (**Fig. A4, B**), which may be connected to the effect of bicarbonate on pyrite oxidation we observed previously (**Fig. A3**).

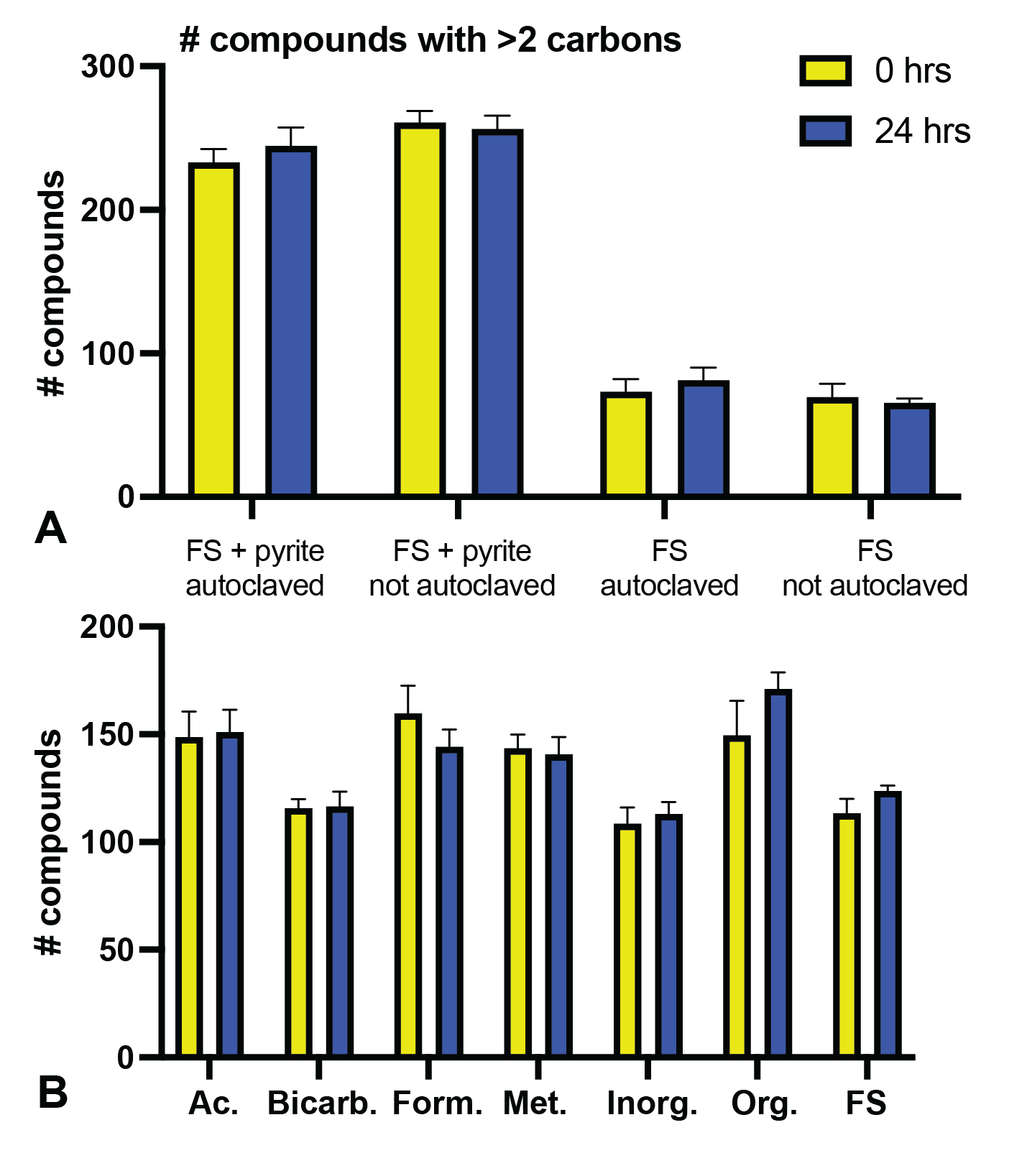

**Fig. A4**. Change in number of detected compounds with more than two carbon atoms with and without pyrite or autoclaving (A) and in autoclaved samples with pyrite with different composition (B): Ac. – only acetic acid; Bicarb. – only sodium bicarbonate; Form. – only formic acid; Met. – only methanol; Inorg. – only inorganic FS components (ammonia, bicarbonate, TMP); Org. – only organic FS components (formic, acetic acids and methanol); FS – complete food set. Error bars are SEM.

**Formation of urea and nitrogen-containing organics.** The only nitrogen-containing compound in FS is ammonia, however, in LCMS data we detect a plethora of various nitrogen-containing organics. Additionally, the experiments were performed under a 95% N_2_ atmosphere. Slow pyrite-catalyzed nitrogen fixation – formation of ammonia sulfate from molecular nitrogen was shown to occur even without UV photocatalysis (Mateo-Marti et al., 2019).

A notable nitrogen-containing compound consistently detected in FS is urea, and it may be the source of some of the nitrogen-containing compound diversity in our samples. Urea synthesis from ammonia and CO_2_ is possible via the Basarov reaction, however it typically requires extreme conditions to achieve high yields (Meessen, 2014). Alternative source of nitrogen organics in our samples could be cyanide (Das et al., 2019), however it is difficult to prove with our experimental setup. We detect the most urea (**Fig. A5, A**) and other nitrogen-containing compounds (**Fig. A7**) in the presence of pyrite, implying possibility of pyrite catalysis. Similarly, formation of an amide bond with formic acid resulting in urea-1-carboxylate occurs largely in the presence of pyrite (**Fig. A5, B**).

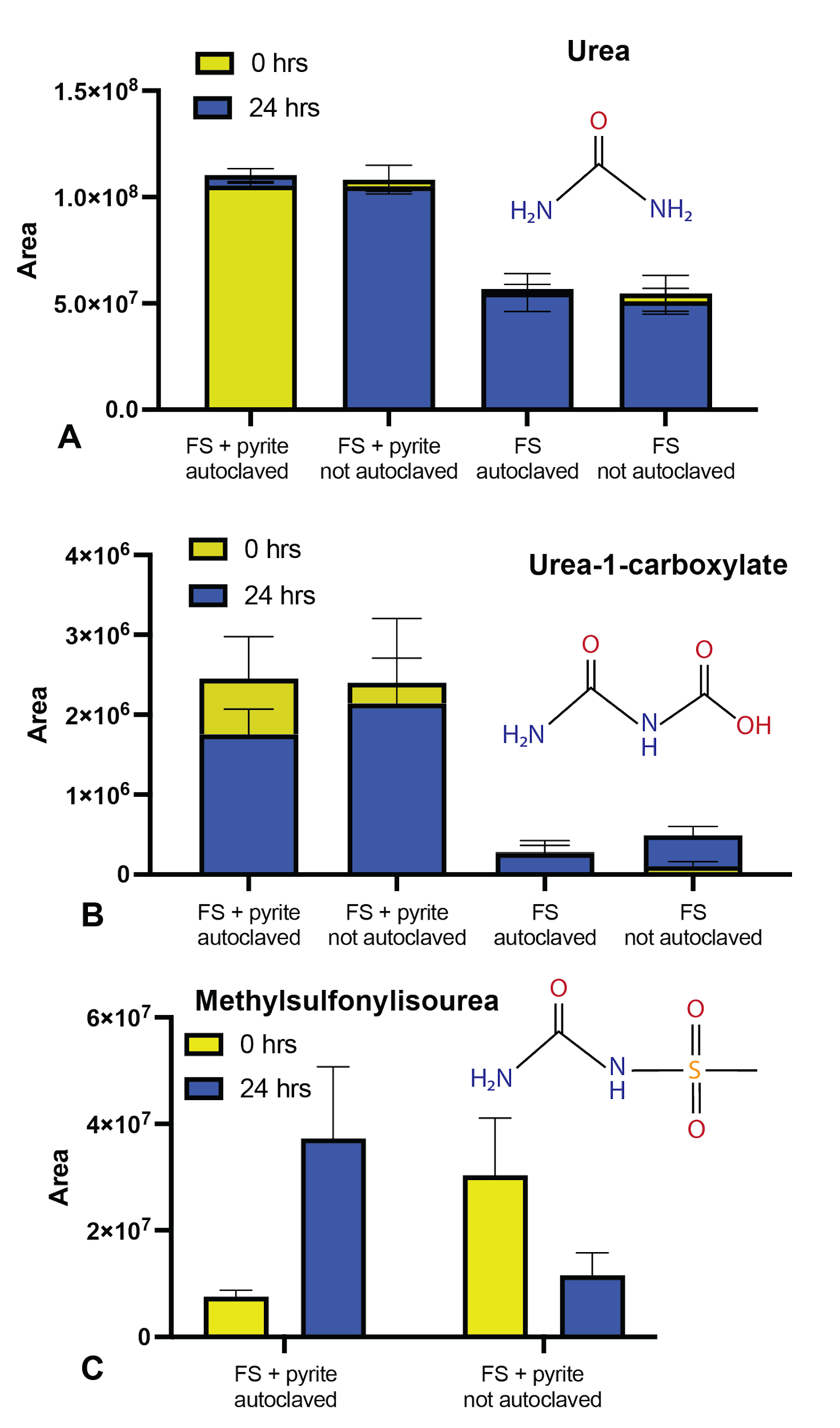

**Fig. A5**. Relative abundance of urea-related products: urea (A), urea-1-carboxylate (B) and methylsulfonylisourea (C) changing due to autoclaving, presence of pyrite and incubation. Error bars are SEM.

We also find significant amounts of urea in pyrite-containing autoclaved samples that only included ammonia (**Fig. A6, A**). Additionally, we find a significant amount of other nitrogen-containing compounds in those samples (**Fig. A6, B**). While there were no explicit carbon sources in the solution, it is likely that trace dissolved carbonate or atmospheric CO_2_ was sufficient for pyrite-catalyzed reactions. In addition to urea, we also detect other nitrogen-containing organic compounds such as amines and heterocycles.

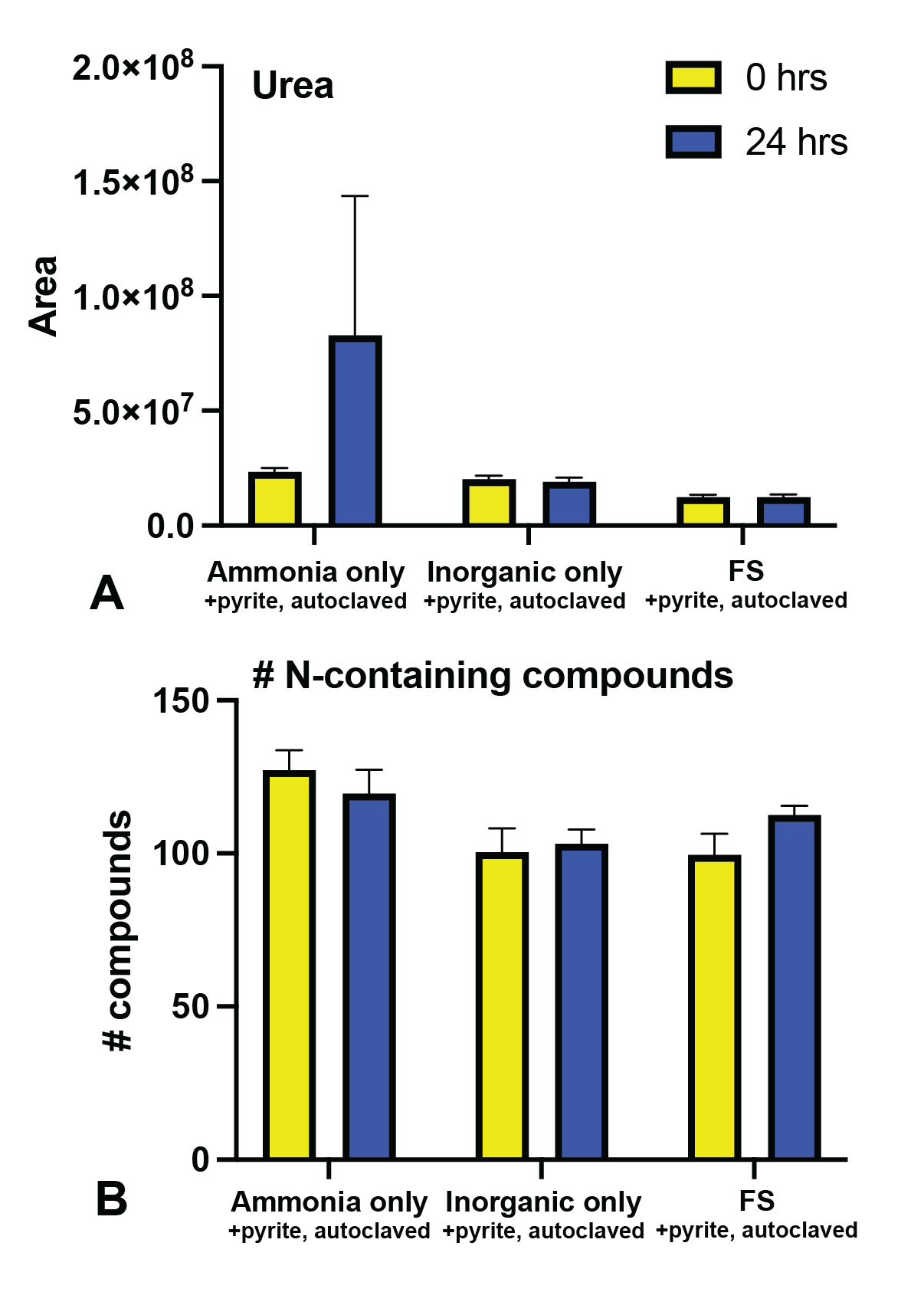

**Fig. A6.** Relative abundance of urea (A), and number of detected nitrogen-containing compounds (B) in autoclaved mixtures with pyrite with ammonia, inorganic FS components and complete FS. Error bars are SEM.

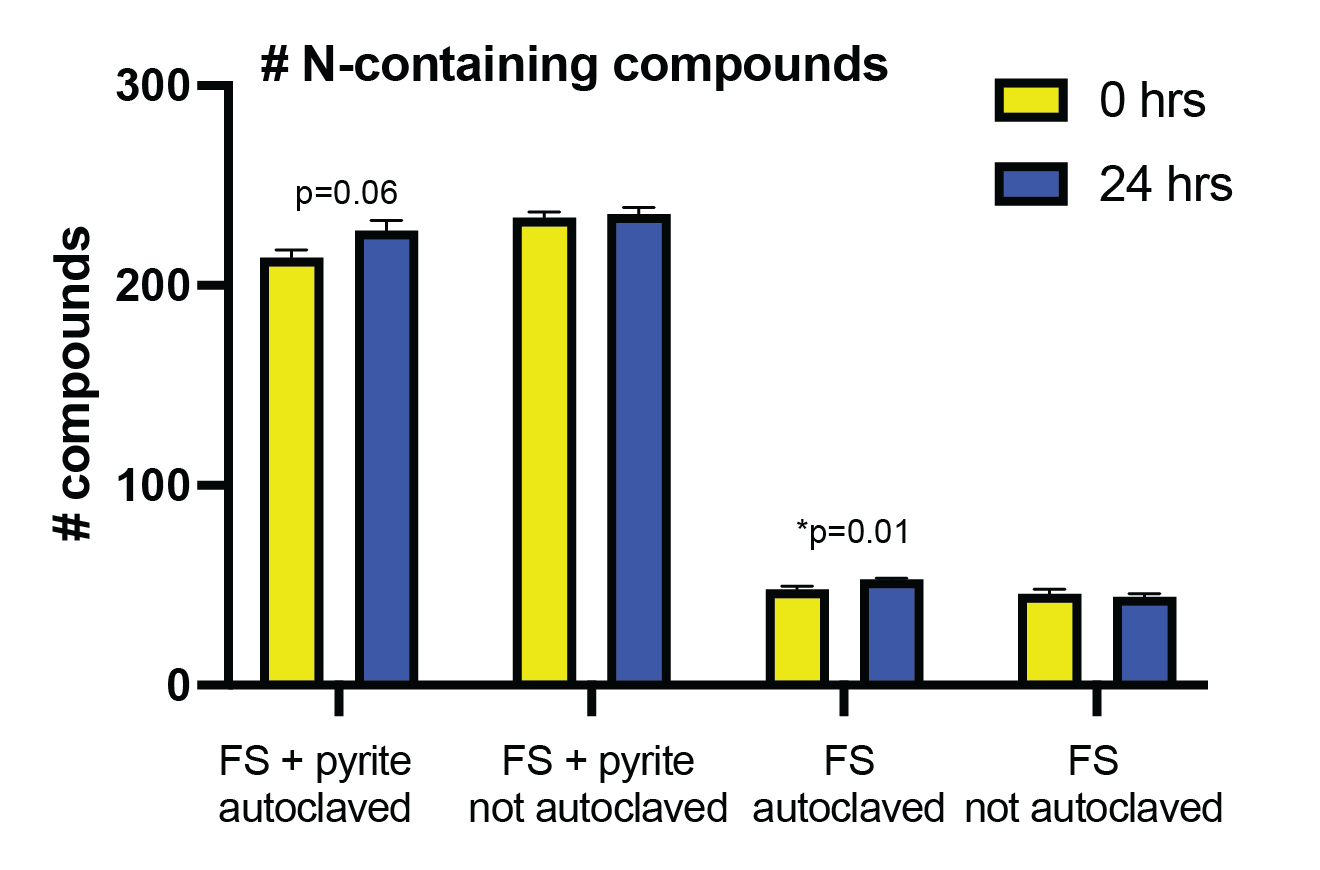

**Fig. A7.** Number of nitrogen-containing compounds in FS with or without pyrite or autoclaving. Error bars are SEM.

**Phosphorylation.** The source of phosphorus in FS is sodium trimetaphosphate (TMP). In the FS solution, TMP seems to convert into linear triphosphate (**Fig. A8, A-B**). We find that pyrite likely catalyzes sequential TMP breakdown into pyrophosphate and then orthophosphate (**Fig. A8, C-D**). This effect could also be enhanced by presence of urea in pyrite-containing samples (**Fig. A5, A**), as urea was shown to promote phosphorylation reactions (Lohrmann & Orgel, 1971)

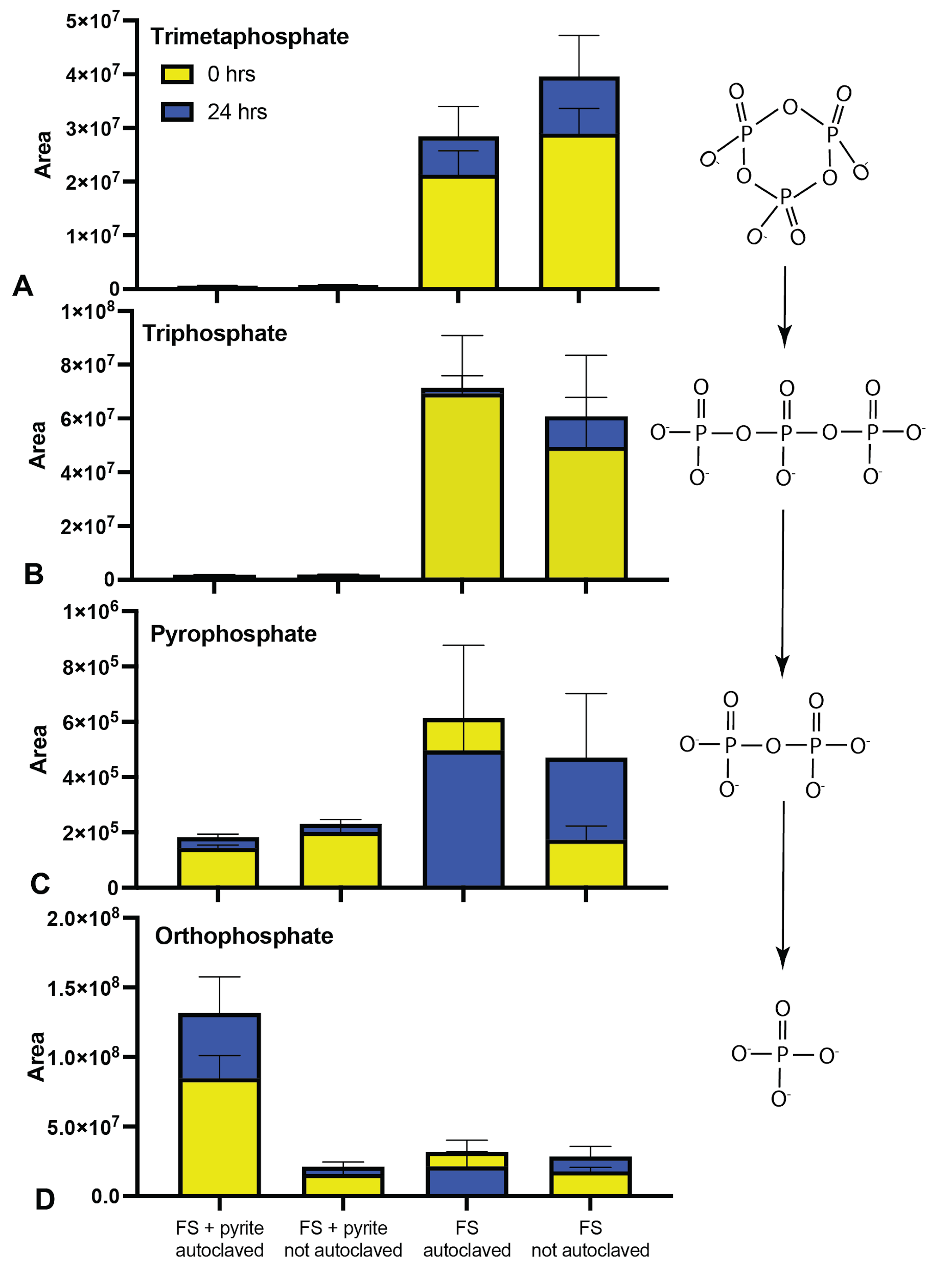

**Fig. A8.** Changes in abundance of various phosphate anions due to presence of pyrite and autoclaving: A – trimetaphosphate; B – triphosphate; C – pyrophosphate; D – orthophosphate. Error bars are SEM.

**Appendix B. Supplementary figures.**

|  | **Analyzed separately with CD** | | | | | | **Analyzed together with CD** | | | | | |
| --- | --- | --- | --- | --- | --- | --- | --- | --- | --- | --- | --- | --- |
|  | FS+P, A. | FS+P, NA. | FS, A. | FS, NA. | W+P, A. | W+P, NA. | FS+P, A. | FS+P, NA. | FS, A. | FS, NA. | W+P, A. | W+P, NA. |
| **Total features** | 1187 | 1156 | 842 | 452 | 919 | 1132 | 4025 | 4025 | 4025 | 4025 | 4025 | 4025 |
| **Area >10^5^** | 867.2  (75.7) | 854.2  (67.8) | 501.2  (112.6) | 308.6  (8.9) | 756.6  (9.0) | 834.8  (111.3) | 2024.6  (150.9) | 2058.8  (114.3) | 1150.6  (177.0) | 987.8  (36.2) | 1905.4  (17.1) | 2061.2  (172.1) |
| **Area >10^6^** | 540.2  (32.9) | 523.6  (22.6) | 166.0  (33.2) | 96.2  (6.5) | 493.2  (7.6) | 516.6  (38.3) | 986.2  (54.4) | 1015.4  (31.1) | 400.2  (47.0) | 317.6  (6.7) | 955.6  (18.0) | 1018  (58.4) |
| **Area >10^7^** | 219.4  (9.8) | 227.0  (4.0) | 41.2  (6.9) | 24.4  (2.2) | 229.6  (10.9) | 230.4  (8.2) | 345.4  (13.2) | 358.4  (9.9) | 108.4  (7.0) | 90.2  (4.2) | 349.8  (12.9) | 367.2  (13.9) |

**Table B1.** Mean counts of detected features using different Compound Discoverer (CD) analyses – first, all replicates from the same treatment were analyzed together in one CD run, then all replicates from every treatment were ran together to allow for cross-treatment comparisons. All samples were incubated for 24 hours; W – water, P – pyrite, A – autoclaved prior to incubation, NA – not autoclaved prior to incubation. Numbers in parentheses represent standard deviations.

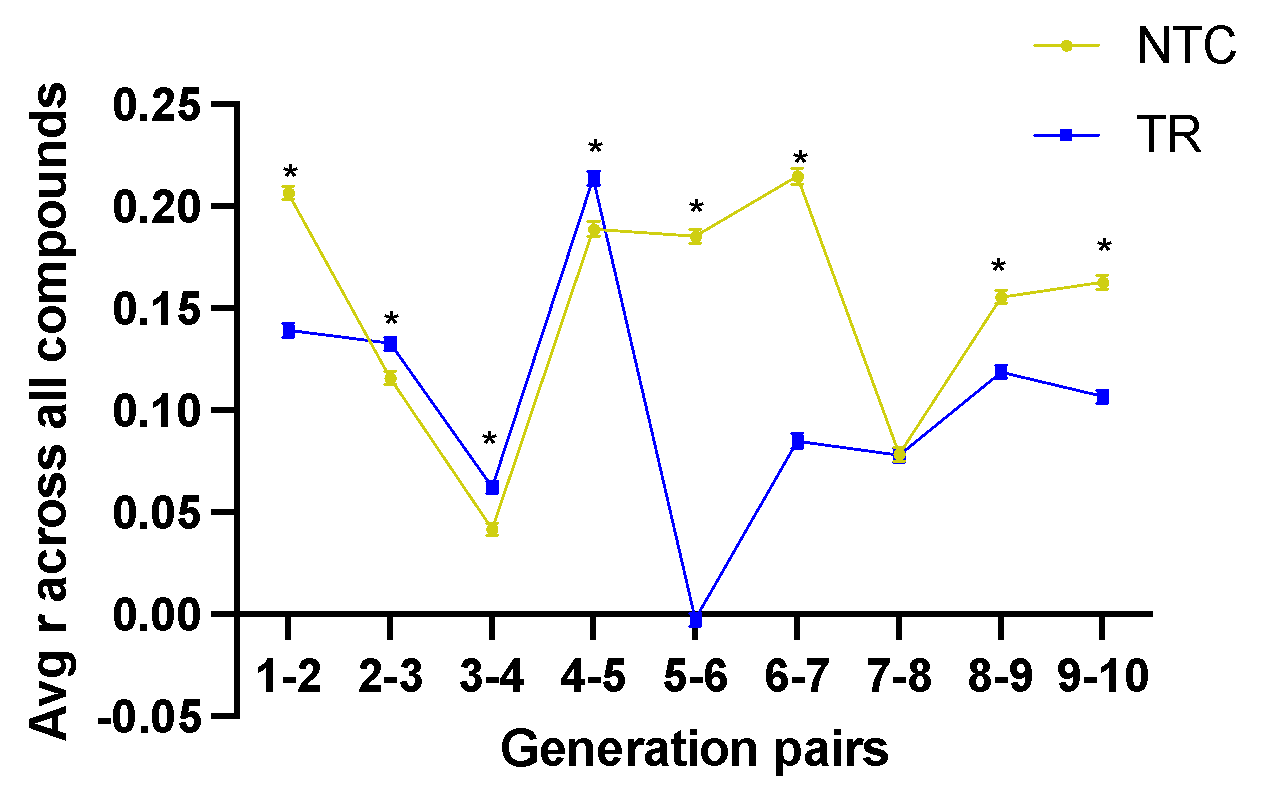

**Fig. B1.** Average Pearson correlation coefficient across all detected compounds in TRs and NTCs.

**Appendix C. Rule-based chemical reaction network design.**

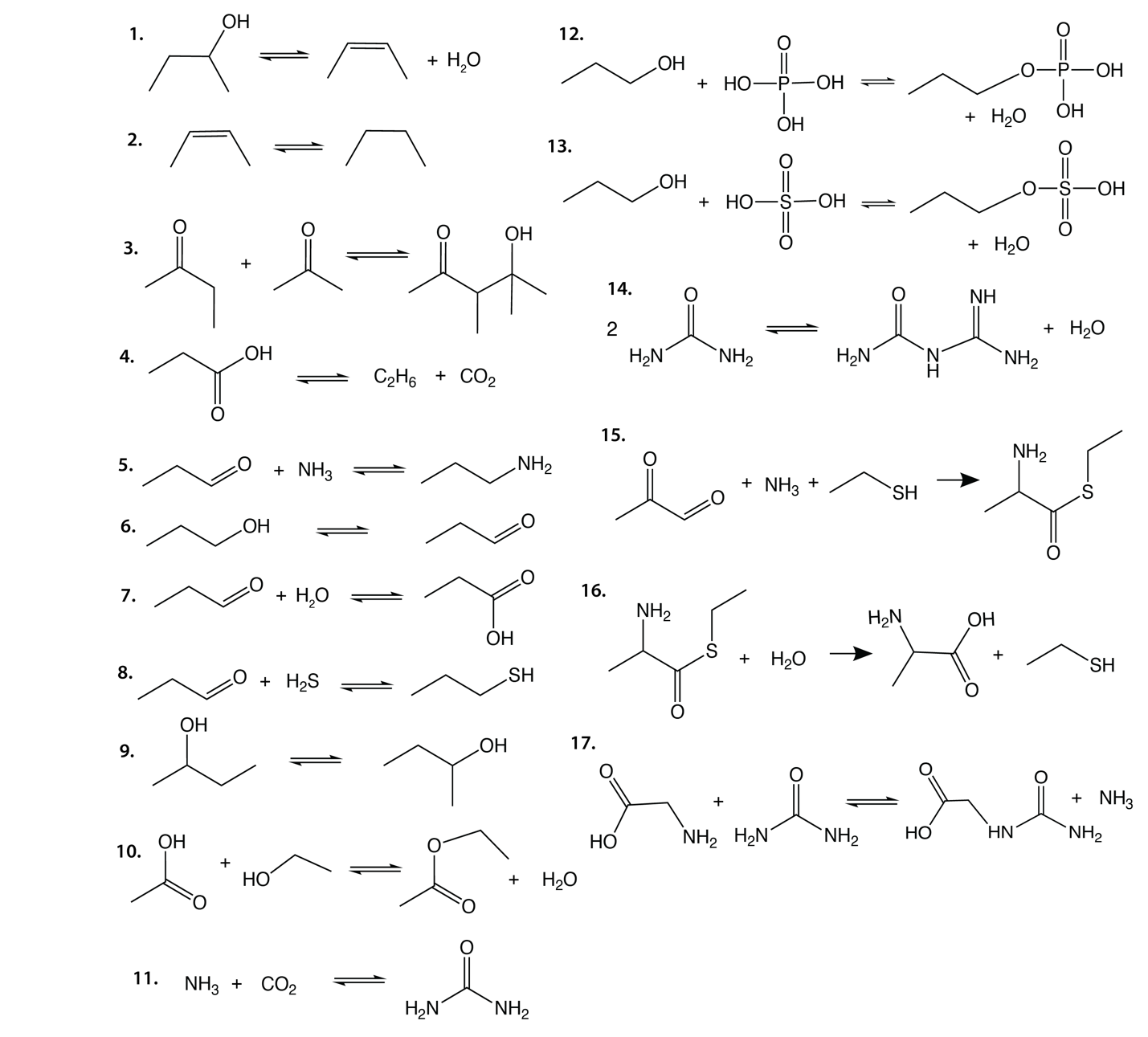

**Fig. C1.** Illustration of selected reaction rules (modified after Fig. 2 in Cuevas-Zuviria & Sokolskyi, 2024).

| **Reaction type** | **Source** |
| --- | --- |
| Dehydration | Muchowska et al., 2019 |
| C=C bond formation | Muchowska et al., 2019 |
| Aldol reaction | Muchowska et al., 2019 |
| Decarboxylation | Muchowska et al., 2019 |
| Iron-catalyzed carbon fixation | Varma et al., 2018 |
| Reductive amination | Muchowska et al., 2019 |
| Amino acid thioesters | Weber, 1998 |
| Thiol synthesis | Kaschke et al., 1994 |
| Urea | Jeilani et al., 2014 |
| Urea-amino acid reactions | Gan et al., 2023 |
| Phosphorylation | Gull, 2014 |
| Sulfonation | Gounaris et al., 2014 |
| Imines | Paczelt et al., 2023 |

**Table C1.** Sources supporting prebiotic plausibility of some of the main classes of reactions in our ruleset.

| **Id** | **Name** | **Reaction SMARTS** |
| --- | --- | --- |
| r001 | Dehydration | [#6:5]-[D2:3]-[#6:2](-[#6:4])-[OH:1]>>[#6:4]\[#6:2]=[#6:3]\[#6:5].[#8:1] |
| r002 | Reduction | [#6:2]=[#6:1]>>[#6:2]-[#6:1] |
| r003 | Aldol reaction | [#6:1]=[O:2].[#6;D2&H2,D1&H3:5]-[#6:4]=[O:3]>>[#8:2]-[#6:1]-[#6:5]-[#6:4]=[O:3] |
| r004 | Neutral decarboxylation | [#6:2]-[#6:1](-[OH:4])=[#8:3]>>[#8:4]=[#6:1]=[#8:3].[#6:2] |
| r005 | Retro-aldol reaction | [OH:2]-[#6:1]-[#6:5]-[#6:4]=[O:3]>>[#6:1]=[O:2].[#6;D2&H2,D1&H3:5]-[#6:4]=[O:3] |
| r006 | Redox decarboxylation | [C:3]-[C:2](=[O:6])-[#6:1](-[OH:4])=[O:5].[OH2:7]>>[O:5]=[C:1]=[O:4].[C:3]-[C:2](-[O:7])=[O:6] |
| r007 | Reductive amination | [CH2:1]=[O:2].[ND0:3]>>[CH2:1]-[#7:3].[#8:2] |
| r008 | Reductive amination | [*;!O:4]~[#6:1]=[O:2].[ND0:3]>>[*;!O:4]-[#6:1]-[#7:3].[#8:2] |
| r009 | Reductive amination | [*:5]-[O:4]-[CH:1]=[O:2].[ND0:3]>>[*:5]-[O:4]-[CH2:1]-[#7:3].[#8:2] |
| r010 | Hydration | [#6:4]\[#6:2]=[#6:3]\[#6:5].[OH2:1]>>[#6:5]-[D2:3]-[#6:2](-[#6:4])-[#8:1] |
| r011 | Alcohol to aldehyde | [CD2,CD1:1]-[OD1:2]>>[#6:1]=[O:2] |
| r012 | Aldehyde to carboxylic acid | [CH2:2]=[O:3].[OD0:4]>>[#6:2](-[O:3])=[O:4] |
| r013 | Aldehyde to carboxylic acid | [*;!O:1]~[CH:2]=[O:3].[OD0:4]>>[*;!O:1]-[C:2](-[O:3])=[O:4] |
| r014 | Aldehyde to carboxylic acid | [*:5]-[O:1]-[CH1:2]=[O:3].[OD0:4]>>[*:5]-[O:1]-[#6:2](-[O:3])=[O:4] |
| r015 | Amino acid thioester synthesis | [C:1]-[C:2](=[O:3])-[CD2:4]=[O:5].[N:6].[SD1:7]-[C:8]>>[C:1]-[C:2]([N:6])-[C:4](=[O:5])-[S:7]-[C:8].[O:3] |
| r016 | AA thioester partial hydrolysis | [C:1]-[C:2]([N:6])-[C:4](=[O:5])-[S:7]-[C:8].[OH2:3]>>[C:1]-[C:2]([N:6])-[C:4](=[O:5])-[O:3].[S:7]-[C:8] |
| r017 | Thiol synthesis from H2S | [C:2]=[O:3].[SD0:4]>>[C:2]-[S:4].[O:3] |
| r018 | Esterification | [#6:1](=[#8:2])-[OH:6].[CH2,CH3:5]-[OH:4]>>[#6:5]-[#8:6]-[#6:1](=[#8:2]).[#8:4] |
| r019 | Esterification w double bond | [#6:1](=[#8:2])-[OH:6].[CH2,CH1:7]=[CH:5]-[OH:4]>>[CH2,CH1:7]=[#6:5]-[#8:6]-[#6:1](=[#8:2]).[#8:4] |
| r020 | Isomerization (hydroxyls) | [#6:2]-[#6:1]-[OH:3]>>[#6:1]-[#6:2]-[OH:3] |
| r021 | Urea condensation | [#7:3]-[#6:2](-[#7:1])=[O:4].[#7:7]-[#6:6](-[#7:5])=[O:8]>>[#7:7]-[#6:6](=[#7:5])-[#7:3]-[#6:2](-[#7:1])=[O:4].[#8:8] |
| r022 | Amino acid/urea condensation | [#7:5]-[#6:6]-[#6:7](-[#8:9])=[O:8].[#7:1]-[#6:2](-[#7:3])=[O:4]>>[#7:3]-[#6:2](=[O:4])-[#7:1]-[#6:6]-[#6:7](-[#8:9])=[O:8].[#7:5] |
| r023 | Ketone synthesis | [#6:1]-[#6:2](-[OH:3])-[#6:4]>>[#6:1]-[#6:2](=[O:3])-[#6:4] |
| r024 | Phosphorylation | [CH2,CH3:1]-[OH:2].[$([P:3]([OD1])([OD1])([OD1])=[OD1])]>>[C:1][O:2][P:3](O)(O)=O.[O] |
| r025 | Organosulfates | [CH2,CH3:1]-[OH:2].[OH:6][S:3]([OH:7])(=[O:4])=[O:5]>>[#6:1]-[#8:7][S:3]([#8:6])(=[O:4])=[O:5].[#8:2] |
| r026 | Iron-catalyzed carbon fixation | [CH1,CH2,CH3,CH4:1].[#8:4]=[#6:2]=[#8:3]>>[#6:1]-[#6:2]=[#8:4].[#8:3] |
| r027 | Aldehyde to alcohol | [CH2:1]=[O:2]>>[CH3:1]-[OH:2] |
| r028 | Aldehyde to alcohol | [*;!O:3]~[CH:1]=[O:2]>>[*;!O:3]-[CH2:1]-[OH:2] |
| r029 | Aldehyde to alcohol | [*:4]-[O:3]-[CH:1]=[O:2]>>[*:4]-[O:3]-[CH2:1]-[OH:2] |
| r030 | Carboxylic acid to aldehyde | [#6:2](-[OH:3])=[O:4]>>[CD2,CD1:2]=[O:3].[OD0:4] |
| r031 | Ketone to alcohol | [#6:1]-[#6:2](=[O:3])-[#6:4]>>[#6:1]-[#6:2](-[OH:3])-[#6:4] |
| r032 | Urea synthesis | [O:3]=[C:2]=[O:1].[ND0:4].[ND0:5]>>[O:1]=[C:2](-[N:4])[N:5].[O:3] |
| r033 | Amination reverse | [CH2,CH3:1]-[NH2:2].[OH2:3]>>[#6:1]=[#8:3].[NH3:2] |
| r034 | Thiols reverse | [CH2,CH3:1]-[SH:2].[OH2:3]>>[#6:1]=[#8:3].[SH2:2] |
| r035 | Phosphorylation reverse | [C:1][O:2][P:3]([OH:4])([OH:5])=[O:6].[OH2:7]>>[CH2,CH3:1]-[OH:2].[OH:7][P:3]([OH:4])([OH:5])=[O:6] |
| r036 | Organosulfates reverse | [C:1][O:2][S:3]([OH:4])(=[O:5])=[O:6].[OH2:7]>>[CH2,CH3:1]-[OH:2].[OH:7][S:3]([OH:4])(=[O:5])=[O:6] |
| r037 | Urea condensation reverse | [#7:7]-[#6:6](=[#7:5])-[#7:3]-[#6:2](-[#7:1])=[O:4].[OH2:8]>>[#7:3]-[#6:2](-[#7:1])=[O:4].[#7:7]-[#6:6](-[#7:5])=[O:8] |
| r038 | Ester hydrolysis | [O:1]=[#6:2]-[O:3]-[#6:4].[OH2:5]>>[O:1]=[#6:2]-[OH:3].[#6:4]-[OH:5] |
| r039 | Amino acid/urea condensation reverse | [#7:3]-[#6:2](=[O:4])-[#7:1]-[#6:6]-[#6:7](-[#8:9])=[O:8].[NX3:5]>>[#7:5]-[#6:6]-[#6:7](-[#8:9])=[O:8].[#7:1]-[#6:2](-[#7:3])=[O:4] |
| r040 | Sulfonylation | [#6:1]=[#6:2].[OH:5]-[S:6](-[OH:7])(=[O:8])=[O:9]>>[#6:1]-[#6:2](-[S:6](-[OH:7])(=[O:8])=[O:9]).[OH2:5] |
| r041 | Sulfonyl hydrolysis | [#6:1]-[#6:2](-[S:6](-[OH:7])(=[O:8])=[O:9]).[OH2:5]>>[#6:1]=[#6:2].[OH:5]-[S:6](-[OH:7])(=[O:8])=[O:9] |
| r042 | Imine synthesis | [CH2:1]=[#8:2].[#6:3]-[NH2:4]>>[#6:1]=[#7:4]-[#6:3].[OH2:2] |
| r043 | Imine synthesis | [*;!O:5]~[#6:1]=[#8:2].[#6:3]-[NH2:4]>>[*;!O:5]~[#6:1]=[#7:4]-[#6:3].[OH2:2] |
| r044 | Imine synthesis | [*:6]-[O:5]-[CH:1]=[#8:2].[#6:3]-[NH2:4]>>[*:6]-[O:5]-[CH:1]=[#7:4]-[#6:3].[OH2:2] |
| r045 | Imine synthesis | [*;!O:5]~[#6:1]=[#8:2].[NH3:4]>>[*;!O:5]~[#6:1]=[#7:4].[OH2:2] |
| r046 | Imine synthesis | [*:6]-[O:5]-[CH:1]=[#8:2].[NH3:4]>>[*:6]-[O:5]-[CH:1]=[#7:4].[OH2:2] |
| r047 | Imine synthesis | [CH2:1]=[#8:2].[NH3:4]>>[#6:1]=[#7:4].[OH2:2] |
| r048 | Imine hydrolysis | [#6:1]=[#7:4]-[#6:3].[OH2:2]>>[#6:1]=[#8:2].[#6:3]-[NH2:4] |
| r049 | C=C bond formation | [#6:2]-[#6:1]>>[#6:2]=[#6:1] |

**Table C2.** Complete ruleset used in this study. Some rules are split into multiple rows to accommodate for various edge cases. Forward and reverse reactions are written as separate rules.

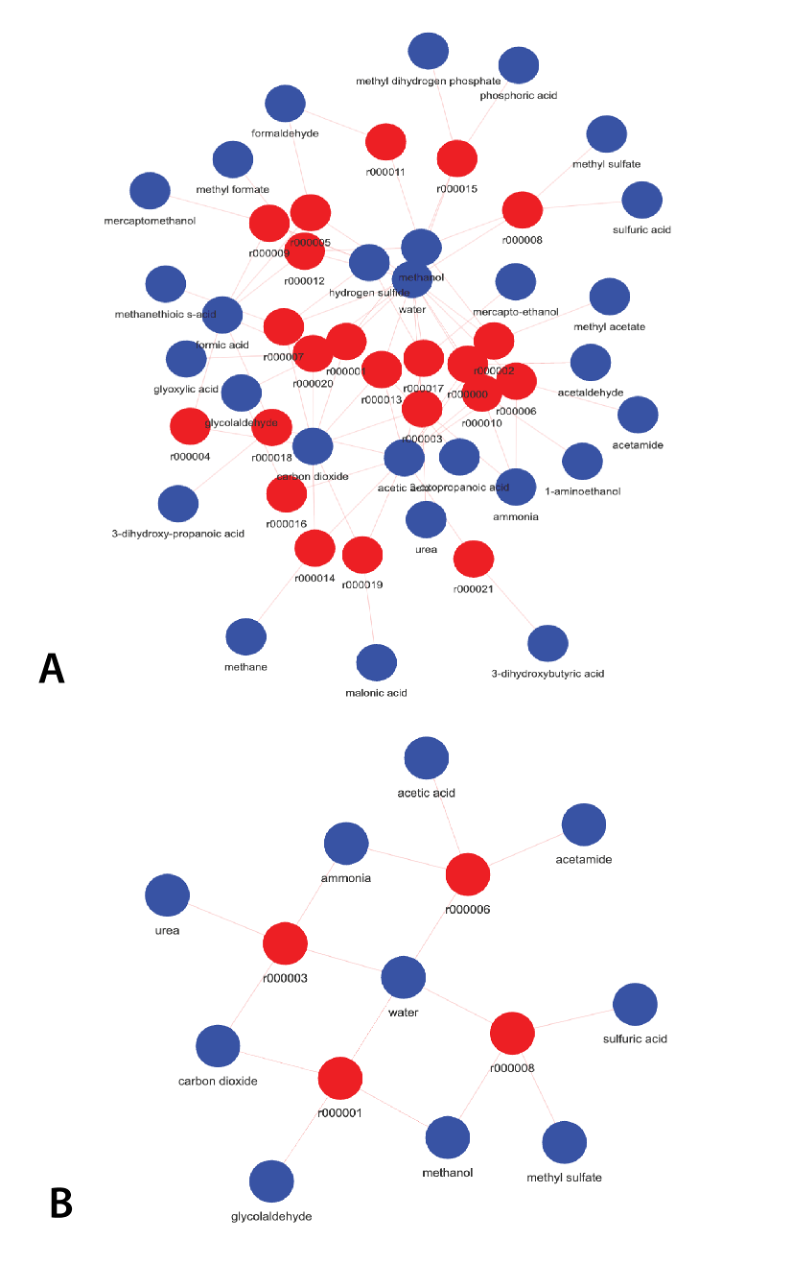

**Fig. C2.** Rule-based CRN resulting from a 1-iteration expansion from the FS using the rules from Table C2: **A** - complete network; **B** - network pruned with compounds detected by LCMS.

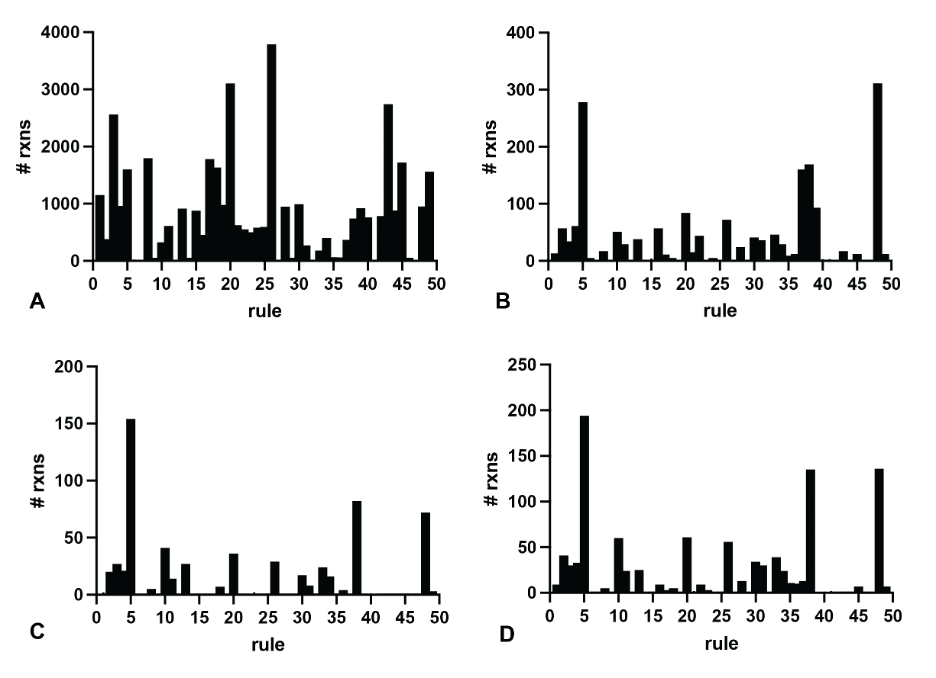

**Fig. C3.** Total number of reactions per rule for the network after 4 iterations of expansion – complete network (**A**), pruned with all compounds detected with LCMS (**B**), just the heritable core (**C**) and just the control core (**D**). Rule numbering is the same as in **Table C2**.
